# ABHD11 inhibition drives sterol metabolism to modulate T cell effector function and alleviate autoimmunity

**DOI:** 10.1101/2025.03.19.643996

**Authors:** Benjamin J. Jenkins, Yasmin R. Jenkins, Fernando M. Ponce-Garcia, Chloe Moscrop, Iain A. Perry, Matthew D. Hitchings, Alejandro H. Uribe, Federico Bernuzzi, Simon Eastham, James G. Cronin, Ardena Berisha, Alexandra Howell, Joanne Davies, Julianna Blagih, Douglas J. Veale, Luke C. Davies, Micah Niphakis, David K. Finlay, Linda V. Sinclair, Benjamin F. Cravatt, Andrew E. Hogan, James A. Nathan, Ursula Fearon, David Sumpton, Johan Vande Voorde, Goncalo Dias do Vale, Jeffrey G. McDonald, Gareth W. Jones, James A. Pearson, Emma E. Vincent, Nicholas Jones

## Abstract

Chronic inflammation in autoimmunity is driven by T cell hyperactivation. This unregulated response to self is fuelled by heightened metabolic programmes, which offers a promising new direction to uncover novel treatment strategies. α/β-hydrolase domain-containing protein 11 (ABHD11) is a mitochondrial hydrolase that maintains the catalytic function of α-ketoglutarate dehydrogenase (α-KGDH), and its expression in CD4+ T cells has been linked to remission status in rheumatoid arthritis (RA). However, the importance of ABHD11 in regulating T cell metabolism and function – and thus, the downstream implication for autoimmunity – is yet to be explored. Here, we show that pharmacological inhibition of ABHD11 dampens cytokine production by human and mouse T cells. Mechanistically, the anti-inflammatory effects of ABHD11 inhibition are attributed to increased 24,25-epoxycholesterol (24,25-EC) biosynthesis and subsequent liver X receptor (LXR) activation, which arise from a compromised TCA cycle. The impaired cytokine profile established by ABHD11 inhibition is extended to two patient cohorts of autoimmunity. Importantly, using a murine model of accelerated type 1 diabetes (T1D), we show that targeting ABHD11 suppresses cytokine production in antigen-specific T cells and delays the onset of diabetes *in vivo*. Collectively, our work provides pre-clinical evidence that ABHD11 is an encouraging drug target in T cell-mediated autoimmunity.

**Graphical Abstract:** 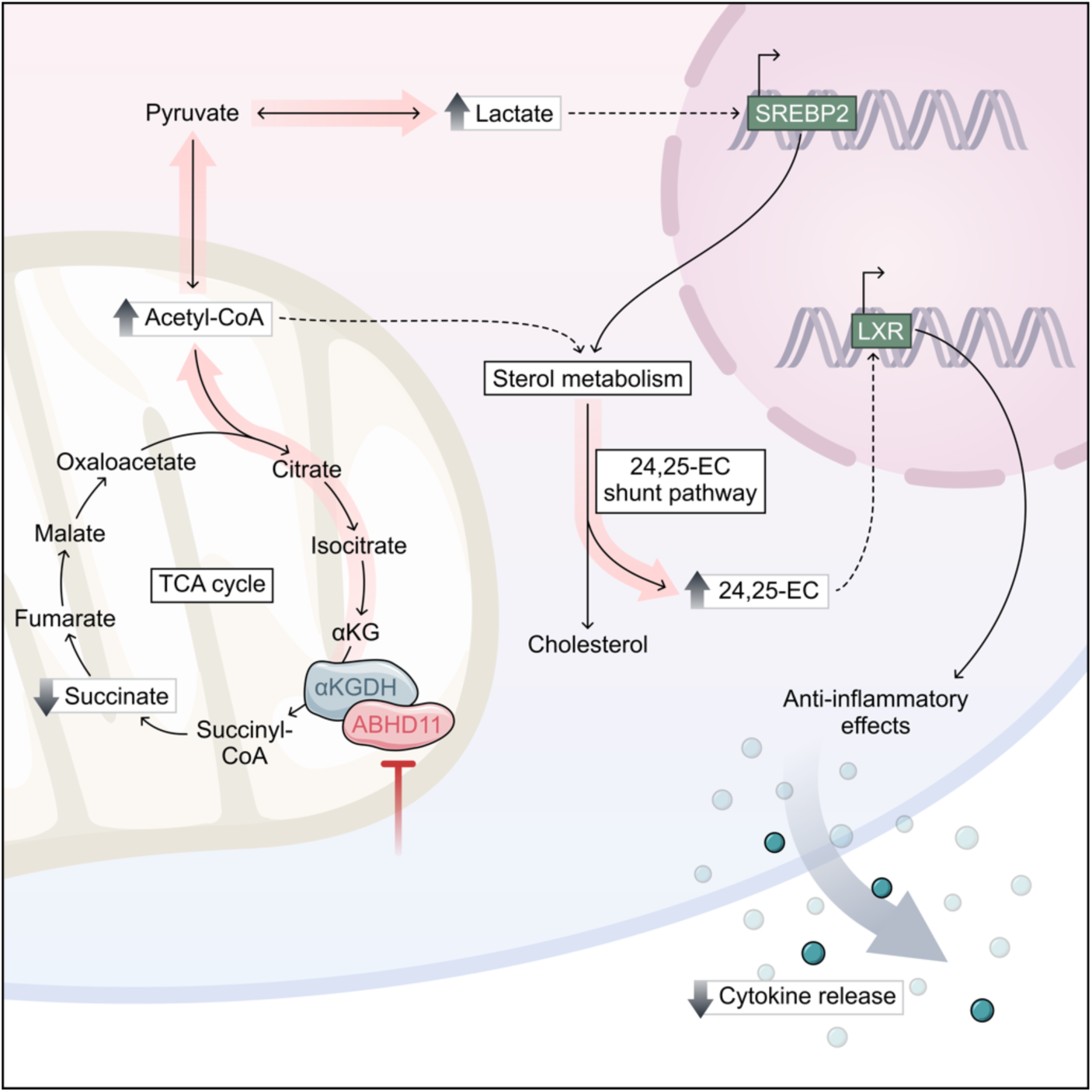

## Introduction

Activated T cells undergo extensive metabolic reprogramming to support the biosynthetic and energetic demands upon antigen encounter^1, 2^. This anabolic switch supports effector function, providing the necessary precursors for cytokine production and blastogenesis. Failure to appropriately regulate T cell activation culminates in autoimmunity, whereby augmented T cell function is driven by alternate or heightened metabolic programmes^3, 4^. Increasingly, altered mitochondrial metabolism within the T cell compartment is being implicated in the onset of autoimmunity. For instance, in rheumatoid arthritis (RA), CD4+ T cells lack mitochondrial aspartate, which disrupts the regeneration of metabolic cofactors required for ER sensor modification, to become enriched in rough endoplasmic reticulum capable of producing large amounts of TNFα, driving inflammation within surrounding tissues^5^. This arthritogenic phenotype arises from mitochondrial malfunction, whereby mitochondrial DNA damage uncouples oxidative phosphorylation (OXPHOS) from ATP production^6^, whilst defective succinyl-CoA ligase function drives reductive carboxylation^7^. Indeed, TNFα itself is likely to be a major driver of these changes in mitochondrial metabolism^8^. Mitochondrial dysfunction is a feature of many autoimmune conditions, as pathogenic T cells in systemic lupus erythematosus (SLE) can be characterised by elevated levels of OXPHOS^4, 9^, whilst their counterparts in multiple sclerosis (MS) display reduced levels of mitochondrial respiration^10^. These distinct modes of mitochondrial dysfunction highlight the metabolic heterogeneity that underpins autoimmunity.

Several leading immunosuppressants target the immunometabolic profile to modulate inflammation^11^. For example, methotrexate has long been a first-line treatment of many autoimmune disorders, inhibiting key aspects of folate metabolism and nucleotide synthesis to limit immune cell function^12^. However, a substantial proportion of autoimmune patients do not respond to methotrexate^13^, whilst other treatments are often blighted by debilitating side effects^14^. To this end novel treatment strategies have been developed. Both tetramerisation of pyruvate kinase^15^ and inhibition of ATP synthase^16^ have improved disease outcomes in experimental autoimmune encephalomyelitis (EAE) models, harnessing the relationship between metabolism and autoimmunity for therapeutic benefit. Moreover, metabolic modulators such as 2-deoxyglucose and metformin have shown promise alongside traditional treatment strategies in lupus-prone mice^17^, which, amongst other studies, has catalysed the repurposing of metformin in SLE^18^, RA^19^ and multiple sclerosis^20^ in clinical trials. Thus, there is substantial precedent for manipulating T cell metabolism to uncover alternative therapeutic agents that alleviate autoimmunity.

Molecular profiling has identified α/β-hydrolase domain-containing protein 11 (ABHD11) as one of the genes most associated with remission status in RA^21^. Specifically, reduced ABHD11 expression within patient-derived CD4+ T cells correlates with improved disease outcome and clinical remission^21^. AHBD11 plays a peripheral role in mitochondrial metabolism, where it regulates the activity of α-ketoglutarate dehydrogenase (α-KGDH), the rate-limiting enzyme within the TCA cycle, by maintaining functional lipoylation of its catalytic subunit, dihydrolipoamide S-succinyltransferase (DLST)^22^. In the absence of ABHD11 function, α-ketoglutarate (α-KG) accumulates and is subsequently converted to 2-hydroxyglutarate (2-HG), which can inhibit the function of α-KG-dependent dioxygenases involved in epigenetic remodelling and hypoxia-inducible factor (HIF) signalling^22^. Whilst 2-HG has previously been shown to suppress cytokine production and cytotoxicity in murine CD8+ T cells^23^, the importance of ABHD11 function in human T cells has yet to be determined.

Here, we show that loss of ABHD11 function limits T cell cytokine production. Mechanistically, we demonstrate these changes are not attributed to a 2-HG epigenetic axis, rather an augmented oxysterol biosynthesis programme that arises following the accumulation of lactate and acetyl-CoA. To this end, heightened intracellular lactate promotes SREBP2 signalling, which upregulates sterol biosynthetic processes to drive the production of 24,25-epoxycholesterol (24,25-EC) via a mevalonate shunt pathway and subsequent activation of liver X receptor (LXR) signalling. Crucially, *ex vivo* pharmacological inhibition of ABHD11 suppressed T cell function in two autoimmune patient cohorts, whilst also attenuating disease activity in a murine model of autoimmunity *in vivo*. Together, these data demonstrate the potential of modulating ABHD11 function for therapeutic benefit in T cell-mediated autoimmune disease.

## Results

### ABHD11 is required for murine and human T cell effector function

Given that reduced expression of ABHD11 within CD4+ T cells is associated with remission in RA^21^, we sought to better understand the role of ABHD11 in T cell biology. Firstly, we established the degree of ABHD11 protein expression in human CD4+ T cells, wherein ABHD11 was expressed at low levels in unstimulated T cells, but became notably upregulated upon TCR-mediated activation (Figure 1A). To determine the importance of ABHD11 function, human CD4+ T cells were treated with ML-226, a highly-selective inhibitor that targets the active site serine of ABHD11^24^, whilst concomitantly being activated by anti-CD3 and anti-CD28 for 24 h. Here, ML-226 significantly impaired cytokine production, with striking reductions in the release of IL-2, IL-10, IL-17, IFNγ and TNFα (Figure 1B). We also analysed underlying mRNA levels for *IL17* and *IFNG*, where comparable reductions were observed at the gene transcript level (Figure 1C). Although there were modest reductions in CD25, CD44 and CD69 expression following ABHD11 inhibition, the proportion of cells that upregulate the expression of these markers following TCR stimulation remained similar (Figure 1D; Supplementary Figure 1A), indicating that early TCR signalling events are intact. Despite the loss of effector function, ABHD11 inhibition did not induce any notable reduction in T cell size (Figure 1E), further demonstrating that early activation signals are unperturbed. In line with this, global protein translation, as measured by puromycin incorporation, was not significantly altered by ABHD11 inhibition (Supplementary Figure 1B). Importantly, cell viability remained intact in T cells exposed to ML-226 (Supplementary Figure 1C), confirming that the observed loss of effector function is attributable to ABHD11 inhibition rather than compromised cell viability.

**Figure 1.**
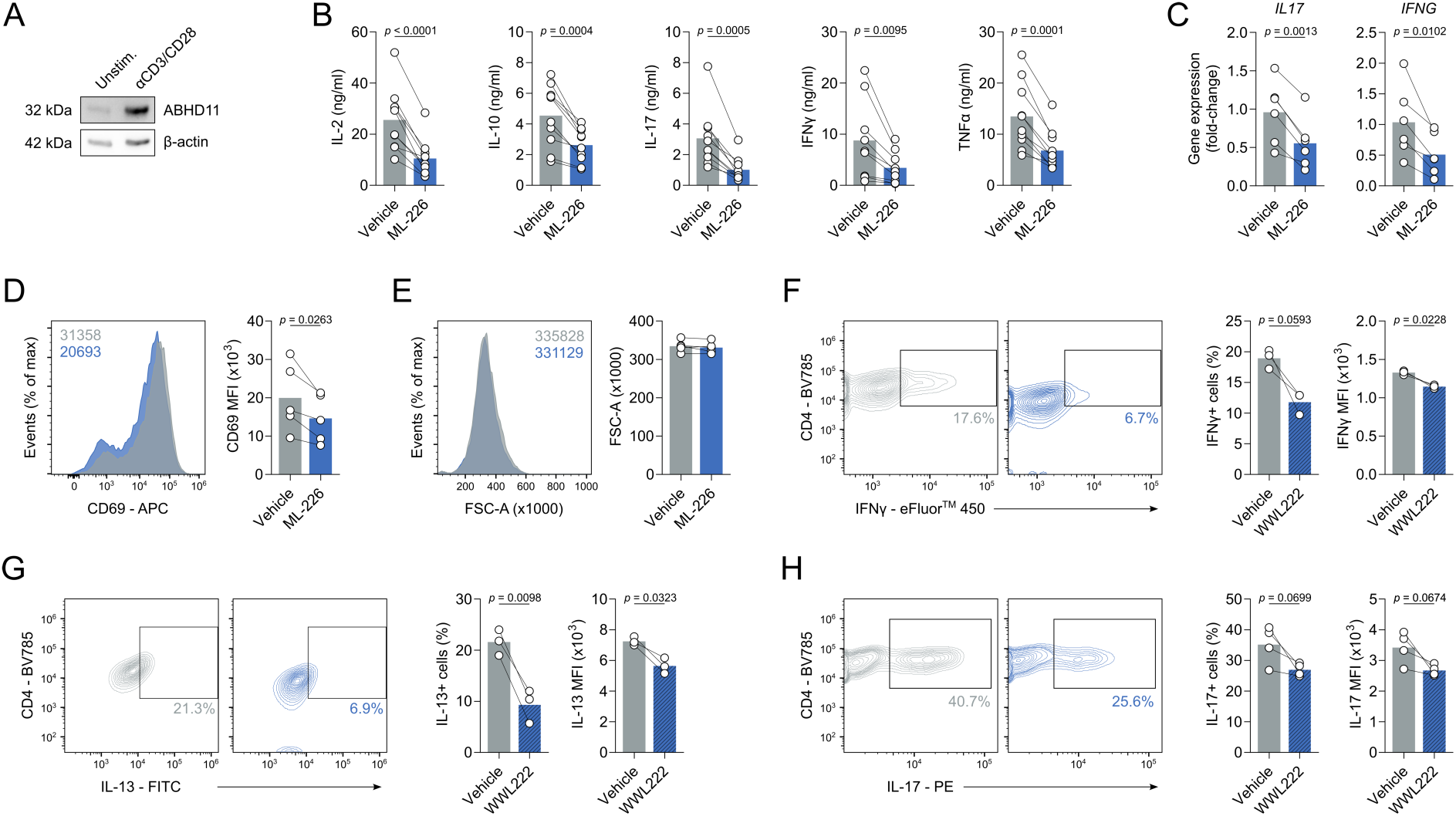
ABHD11 inhibition impairs T cell activation and cytokine production. (**A**) ABHD11 expression in CD4+ T cells, unstimulated or activated with α-CD3 and α-CD28 (n = 3). Protein loading assessed using β-actin. (**B**) IL-2, IL-10, IL-17, IFNγ and TNFα production by CD4+ effector T cells (n = 9/10). (**C**) qPCR analysis of *IL17* and *IFNG* expression in CD4+ effector T cells (n = 6). (**D**) Surface expression of CD69 as a measure of activation, as measured by flow cytometry, on CD4+ effector T cells (n = 5). (**E**) Cell size, as determined by forward scatter area, of CD4+ effector T cells (n = 5). (**F**) Intracellular IFNγ expression, as measured by flow cytometry, in murine CD4+ effector T cells following polarisation towards Th1 in the presence and absence of WWL222 (n = 3). (**G**) Intracellular IL-13 expression, as measured by flow cytometry, in murine CD4+ effector T cells following polarisation towards Th2 in the presence and absence of WWL222 (n = 3). (**H**) Intracellular IL-17 expression, as measured by flow cytometry, in murine CD4+ effector T cells following polarisation towards Th17 in the presence and absence of WWL222 (n = 3). Experiments were carried out using human samples, unless otherwise stated. CD4+ T cells were activated with α-CD3 and α-CD28 for 24 h, in the presence and absence of ML-226, unless otherwise stated. Data are expressed as mean, with paired dots representing biological replicates.

To further investigate the impact of ABHD11 function on T cell fitness, we activated murine T cells in the presence and absence of WWL222, a potent and selective inhibitor targeting ABHD11 in mice^25^. WWL222 is structurally distinct from ML-226, containing a different chemical scaffold, and, importantly, targets an alternate region of ABHD11^24^. Specifically, murine T cells were polarised *in vitro* towards distinct T cell lineages in the presence and absence of WWL222 to assess its subset-specific effect on cytokine production. As with their human counterparts, cytokine production was impaired by ABHD11 inhibition, with a reduction in both the frequency of IFNγ-, IL-13- and IL-17-producing cells, as well as the quantity of each cytokine, under Th1-, Th2- and Th17-polarising conditions, respectively (Figure 1F-H). Together, these data indicate that ABHD11 is essential for optimal T cell function.

### ABHD11 inhibition rewires human T cell metabolism

Given that ABHD11 primarily maintains the function of α-KGDH within the TCA cycle^22^, we next investigated the impact of ABHD11 inhibition on human T cell metabolism. Firstly, we assessed whether ML-226 directly inhibits the catalytic activity of α-KGDH. Although modest, we recorded a significant reduction in α-KGDH activity (Figure 2A), indicating that ML-226 can inhibit ABHD11 oxidative decarboxylation of α-KG in T cells. Subsequently, we activated T cells in the presence and absence of ML-226 and examined cellular metabolism using a mitochondrial stress assay. Here, there was a significant reduction in oxygen consumption rate (OCR) following ABHD11 inhibition, with consistent reductions in basal respiration, ATP-linked respiration, maximal respiratory capacity and spare respiratory capacity (Figure 2B-C). As expected, these changes precede impaired ATP production from OXPHOS (Figure 2D). Furthermore, we utilised targeted mass spectrometry analysis to understand potential alterations at the metabolite level. Here, ABHD11 inhibition revealed a compromised TCA cycle, in which succinate levels were significantly reduced following ABHD11 inhibition, in addition to a striking accumulation of acetyl-CoA (Figure 2E). Although α-KG levels remain similar following ABHD11 inhibition (Figure 2E), there is a marked increase in the observed α-KG / succinate ratio within these cells (Figure 2F), further indicating impaired α-KGDH activity. However, intracellular 2-HG levels were unchanged (Supplementary Figure 2A), suggesting that there is alternative mechanism underpinning the observed phenotype, rather than the epigenetic mechanism previously described following ABHD11 loss^22^. Moreover, the amino acid pool is also noticeably diminished in these cells, with observed reductions in glutamate, aspartate and asparagine (Figure 2G). Aspartate plays an important role in nucleotide synthesis^26^; therefore, it is unsurprising that there is a concomitant reduction in the production of several nucleotides following ABHD11 inhibition (Supplementary Figure 2B). Interestingly, mitochondria appeared to be larger in T cells treated with ML-226 (Figure 2H), though this translated into no discernible effect on mitochondrial depolarisation (Figure 2I). Equally, despite the reduced glutathione pool present in these cells, this had no effect on mitochondrial reactive oxygen species production (Supplementary Figure 2C-D). Taken together, there is substantial evidence that the TCA cycle is compromised following ABHD11 inhibition, altering the metabolic landscape within the mitochondria.

**Figure 2.**
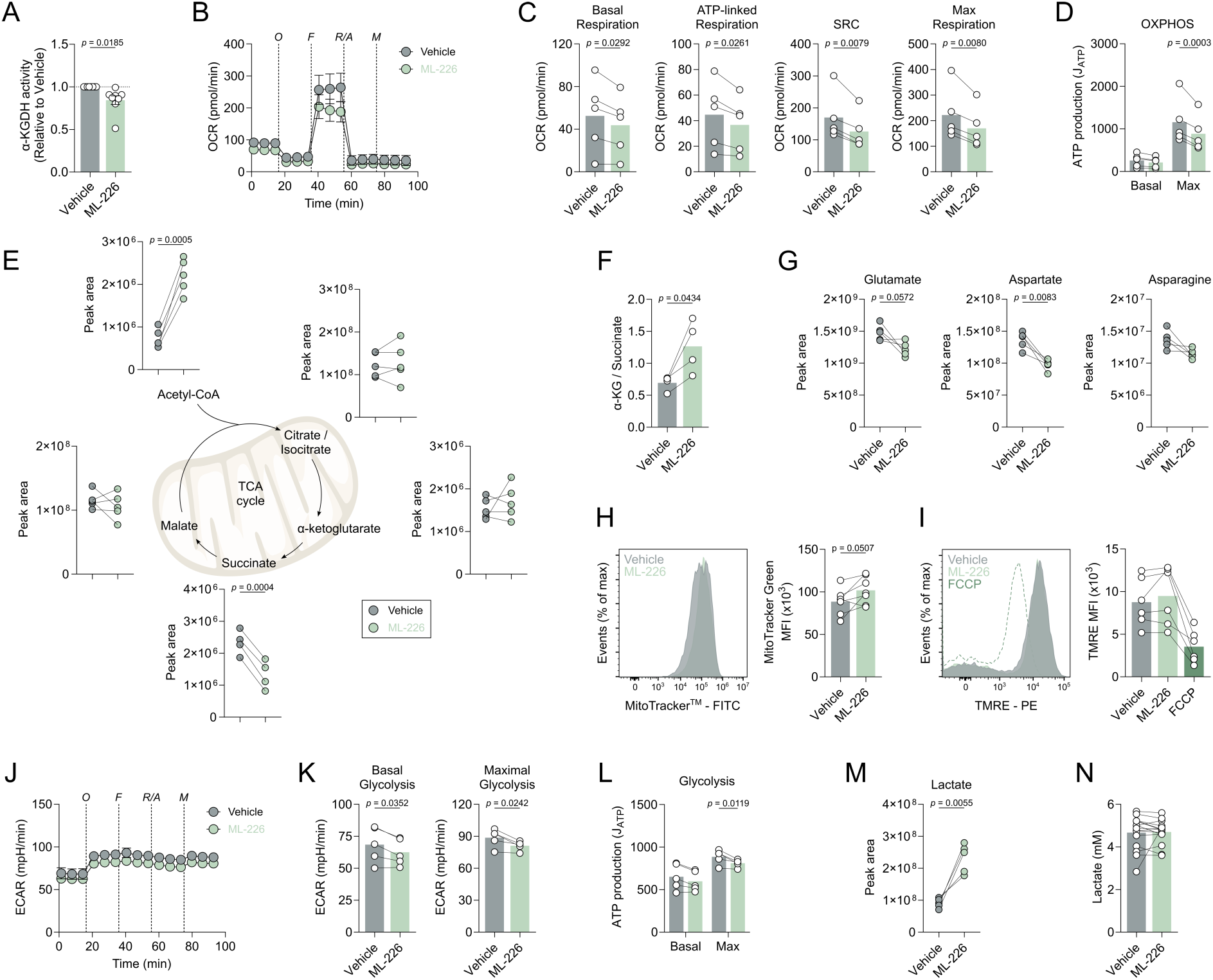
ABHD11 inhibition rewires mitochondrial metabolism in human T cells. (**A**) α-ketoglutarate dehydrogenase (α-KGDH) activity in CD4+ effector T cells (n = 8). (**B**) Oxygen consumption rate (OCR) in CD4+ effector T cells (n = 5). Pre-optimised injections include: oligomycin, FCCP, antimycin A/rotenone (all 1 μM) and monensin (20 μM). (**C**) Calculated OCR parameters including basal respiration, ATP-linked respiration, spare respiratory capacity and maximal respiratory capacity, from OCR measured in (**B**). (**D**) Basal and maximal ATP production (J_ATP_), from OCR measured in (**B**). (**E**) Intracellular levels of selected TCA cycle intermediates in CD4+ effector T cells (n = 5). Metabolites include: acetyl-CoA, citrate/isocitrate, α-ketoglutarate (α-KG), succinate and malate. (**F**) Determination of intracellular α-KG to succinate ratio in CD4+ effector T cells (n = 5). (**G**) Intracellular levels of selected amino acids in CD4+ effector T cells (n = 5). Metabolites include: glutamate, aspartate and asparagine. (**H**) Mitochondrial content, as determined by MitoTracker™ Green, in CD4+ effector T cells (n = 6). (**I**) Mitochondrial membrane potential, as determined by TMRE staining, in CD4+ effector T cells (n = 6). FCCP (1 μM) was used as a positive control. (**J**) Extracellular acidification rate (ECAR) in CD4+ effector T cells (n = 5). Pre-optimised injections include: oligomycin, FCCP, antimycin A/rotenone (all 1 μM) and monensin (20 μM). (**K**) Calculated ECAR parameters including basal glycolysis and maximal glycolysis, from ECAR measured in (**J**). (**L**) Basal and maximal ATP production (J_ATP_), from ECAR measured in (**J**). (**M**) Intracellular levels of lactate in CD4+ effector T cells (n = 5). (**N**) Extracellular lactate in cell-free supernatants from cultures of CD4+ effector T cells (n = 15). All experiments were carried out using human samples. CD4+ T cells were activated with α-CD3 and α-CD28 for 24 h, in the presence and absence of ML-226, unless otherwise stated. Data are expressed as either: mean, with paired dots representing biological replicates; or mean ± SEM.

Given the impact on OXPHOS, we next determined whether ABHD11 inhibition perturbed lactic acid excretion, measured using the extracellular acidification rate (ECAR). We observed a modest reduction in ECAR, whereby glycolytic capacity was attenuated by ABHD11 inhibition (Figure 2J-K). This translated into a reduction in glycolytic ATP production (Figure 2L). However, these changes are not necessarily underpinned by depletion of the metabolite pool, as intracellular levels of glycolytic intermediates are not consistently changed following ABHD11 inhibition (Supplementary Figure 2E). A substantial increase of intracellular lactate levels was observed upon ABHD11 inhibition (Figure 2M). Importantly, the lactate accumulates within these cells, and is not exported at a higher rate (Figure 2N), which agrees with the modest reduction in extracellular acidification rate with ABHD11 inhibition (Figure 2J-L). Together, these data show that ABHD11 inhibition rewires mitochondrial metabolism, establishing a compromised TCA cycle that promotes the accumulation of acetyl-CoA and lactate.

To determine the source of the elevated lactate, we next employed liquid chromatography mass spectrometry to track the fate of cellular metabolites. Here, we performed stable isotope labelling using uniformly-labelled ^13^C_6_-glucose to follow the incorporation of these carbons into downstream metabolites. An elevated percentage of ^13^C is incorporated into lactate (Supplementary Figure 3A), which perhaps suggests a feedback mechanism from the compromised TCA cycle. We also observed ^13^C incorporation into acetyl-CoA. Strikingly, ^13^C incorporation into succinate is unchanged and there was relatively little incorporation of glucose carbon downstream of α-KG, perhaps due to timepoint restrictions. Therefore, we utilised uniformly-labelled ^13^C_5_-glutamine (almost directly upstream of α-KGDH) to measure glutamine anaplerosis (Supplementary Figure 3B). Here, reduced ^13^C incorporation into succinate was more pronounced, whilst incorporation into other TCA cycle intermediates did not appear to be significantly altered (Supplementary Figure 3B). These data would suggest that the accumulation of lactate and most likely acetyl-CoA is sustained by glucose following ABHD11 inhibition.

### ABHD11 inhibition upregulates SREBP signalling to drive oxysterol synthesis and activate LXR

To further our understanding of the mechanisms underpinning rewired T cell metabolism and suppressed effector function following ABHD11 inhibition, we sought to investigate changes in the T cell transcriptome following ML-226 treatment. RNA-Seq analysis revealed 69 genes that were differentially-expressed upon ABHD11 inhibition, of which 38 were upregulated and 31 were downregulated (Figure 3A). To realise the biological relevance of these changes, we performed pathway enrichment analysis to determine which pathways become up- and downregulated upon ABHD11 inhibition. Unsurprisingly, several pathways associated with T cell function, such as *cellular response to cytokine stimulus* and *inflammatory response*, were downregulated following ABHD11 inhibition (Figure 3B), supporting our observed effects on cytokine production (Figure 1B-C). Conversely, the vast majority of the pathways upregulated in response to ABHD11 inhibition were associated with either sterol or fatty acid metabolism, with *sterol biosynthetic process* emerging as the most enriched pathway overall (Figure 3B). Given the multi-faceted roles of acetyl-CoA, we next examined whether its accumulation (Figure 2E) fuels increased histone acetylation. However, there were no observable differences in any of the histone acetylation marks measured following ABHD11 inhibition (Supplementary Figure 4A). In addition, global protein acetylation appeared to be unaffected in these cells (Supplementary Figure 4B). These data would suggest that the immunomodulatory phenotype observed are likely attributed to sterol biosynthetic processes, rather than alternative acetyl-CoA-mediated processes.

**Figure 3.**
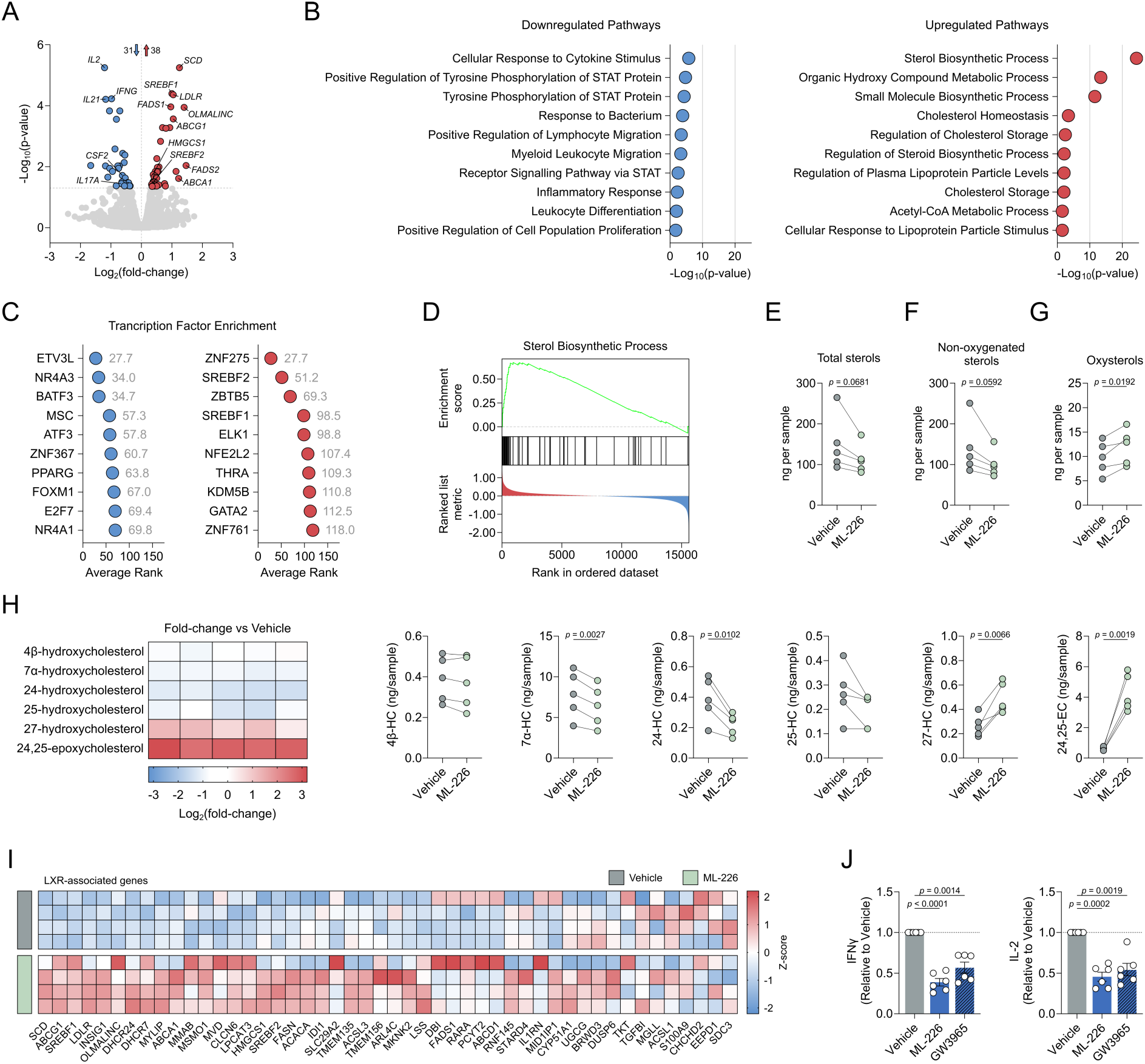
ABHD11 inhibition activates SREBP signalling to drive oxysterol synthesis. (**A**) Differential expression analysis by RNA-Seq in CD4+ effector T cells (n = 4). Blue and red data points represent downregulated and upregulated genes, respectively. Transcripts with an adjusted p-value < 0.05 were considered differentially expressed. (**B**) Pathway enrichment analysis based on differentially-expressed genes. Top 10 enriched pathways are shown. (**C**) Transcription factor enrichment analysis based on differentially-expressed genes. The lower the “Average Rank” value, the more enriched the transcription factor activity. Top 10 enriched transcription factors are shown. (**D**) GSEA enrichment plot for *Sterol Biosynthetic Process*. (**E**) Total intracellular sterol levels in CD4+ T cells (n = 5). (**F**) Intracellular non-oxygenated sterol levels in CD4+ T cells (n = 5). (**G**) Intracellular oxysterol levels in CD4+ T cells (n = 5). (**H**) Intracellular levels of selected oxysterols in CD4+ T cells (n = 5). Metabolites include: 4β-hydroxycholesterol, 7α-hydroxycholesterol, 24-hydroxycholesterol, 25-hydroxycholesterol, 27-hydroxycholesterol, 24,25-epoxycholesterol. Heatmap represented as Log_2_(fold-change) versus vehicle control. (**I**) Expression of LXR-associated genes in CD4+ T cells (n = 4). Heatmap represented as individual gene Z-scores. (**J**) IL-2 and IFNγ production by CD4+ T cells, activated in the presence and absence of ML-226 or GW3965 (n = 6). All experiments were carried out using human samples. CD4+ T cells were activated with α-CD3 and α-CD28 for 24 h, in the presence and absence of ML-226, unless otherwise stated. Data are expressed as either: mean, with paired dots representing biological replicates; or mean ± SEM.

To determine what drives the observed changes in gene expression and, ultimately, the suppressed effector function observed following ABHD11 inhibition, we performed transcription factor analysis. Here, enriched transcription factors are predicted using the differentially-expressed genes identified – the lower the “average rank” score, the more enriched that transcription factor and its activity. Once more, there was a clear association with lipid metabolism, whereby *PPARG*, *SREBF1* and *SREBF2* were all amongst the most enriched transcription factors (Figure 3C). In fact, *SREBF1* and *SREBF2* are significantly upregulated at the transcript level following ABHD11 inhibition (Figure 3A). In further support of this, *sterol biosynthetic process* is the pathway most significantly enriched following ABHD11 inhibition (Figure 3D) – a process tightly regulated by the signalling of sterol regulatory element binding proteins (SREBPs) encoded by *SREBF1* and *SREBF2*^27^. There is recent evidence demonstrating the existence of a lactate-SREBP2 signalling axis within human immune cells^28^. Therefore, given the heightened intracellular lactate levels we observe following ABHD11 inhibition (Figure 2M), we activated T cells in the presence of either ML-226 or lactic acid and assessed whether there was a comparable effect on cytokine production. Here, treatment with lactic acid phenocopied ABHD11 inhibition, with similar reductions in IL-2 and IFNγ production (Supplementary Figure 4C). These data suggest that lactate drives SREBP activation and the subsequent upregulation of sterol biosynthesis pathways.

To establish the significance of augmented sterol biosynthesis following ABHD11 inhibition, we next carried out specialised mass spectrometry to measure the intracellular levels of various sterol species. Surprisingly, we observed a reduction in total sterol levels following ABHD11 inhibition (Figure 3E). This is primarily attributed to a reduction in non-oxygenated sterols, wherein there is a trend towards decrease across the group, with several significantly reduced (Figure 3F; Supplementary Figure 5A). Conversely, we identified heightened oxysterol levels within these cells (Figure 3G), which in turn increased the oxysterol / sterol ratio (Supplementary Figure 5B). Of the oxysterols measured, 27-hydroxycholesterol (27-HC) and 24,25-epoxycholesterol (24,25-EC) emerged as the two most significantly elevated species, particularly 24,25-EC whose levels are approximately 5-10 times higher following ABHD11 inhibition (Figure 3H). Interestingly, there does not appear to be a general increase across all oxysterol species, as 7α-hydroxycholesterol (7α-HC) and 24-hydroxycholesterol (24-HC) levels were reduced upon ABHD11 inhibition (Figure 3H). Intriguingly, we report activation of a shunt pathway – branching from the classical mevalonate pathway at oxidosqualene – which synthesises 24,25-EC from acetyl-CoA (Supplementary Figure 5C).

27-HC and 24,25-EC are potent activators of liver X receptor (LXR) signalling^29^. To establish whether the increased production of 27-HC and 24,25-EC following ABHD11 inhibition leads to LXR activation, we compared treatment with ML-226 versus treatment with GW3965 – a synthetic LXR agonist^30^. Interestingly, recent work by Waddington *et al.* assessed the transcriptional effects of LXR activation on human CD4+ T cell function using RNA-Seq^31^. In comparison to our dataset, 19 of the 65 LXR-regulated transcripts identified were also differentially regulated by AHBD11 inhibition (Figure 3I), which accounts for approximately 27.5% of all differentially-expressed transcripts (Supplementary Figure 5D), indicating significant overlap between both nodes. As such, we hypothesised that LXR activation would phenocopy, at least partially, ABHD11 inhibition in human T cells. To this end, we treated T cells with ML-226 and GW3965 in parallel before assessing cytokine production. Here, we observed comparable reductions in IFNγ and IL-2 production between ABHD11 inhibition and LXR activation (Figure 3J), suggesting that LXR activation drives, at least in part, the anti-inflammatory phenotype observed in human T cells following ABHD11 inhibition.

### ABHD11 inhibition suppresses T cell effector function in autoimmunity

Our findings thus far have outlined that ABHD11 inhibition suppresses T cell effector function, underpinned by a compromised TCA cycle that promotes 24,25-EC synthesis and consequently activates LXR signalling. This highlights the exciting potential of manipulating ABDH11 function for therapeutic benefit in T cell-mediated autoimmune disease, where reversing the hyper-activation and -function of pathogenic T cells would be valuable. We explored this possibility initially in two autoimmune patient cohorts, isolating CD4+ T cells from rheumatoid arthritis (RA) and type 1 diabetes (T1D) patients and activating them in the presence and absence of ML-226 *ex vivo* (Figure 4A). Here, we again observed a significant reduction in the production of a diverse range of cytokines by RA and T1D T cells following ABHD11 inhibition (Figure 4B-C), alongside a modest, but significant reduction in T cell activation (Figure 4D-E). Again, there was a minimal reduction in cell size in both patient cohorts (Supplementary Figure 6A-B), whilst impaired effector function was independent of any changes in viability (Supplementary Figure 6C-D).

**Figure 4.**
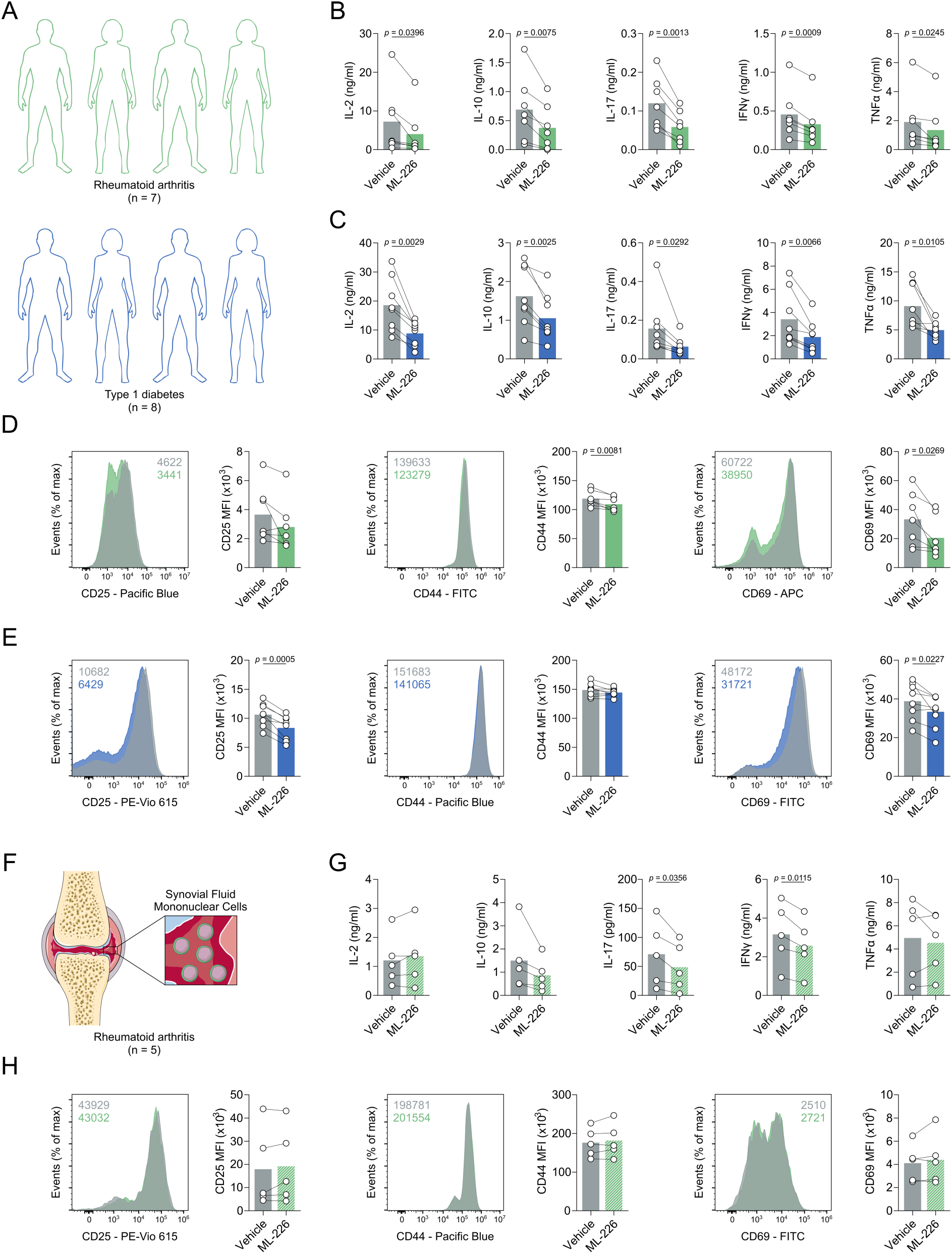
ABHD11 inhibition impairs CD4+ T cell function in autoimmunity. (**A**) Experimental design of autoimmune cohort (rheumatoid arthritis [RA] and type 1 diabetes) CD4+ T cells. (**B-C**) IL-2, IL-10, IL-17, IFNγ and TNFα production by patient-derived CD4+ T cells in autoimmune cohorts of (**B**) RA (n = 7) and (**C**) T1D (n = 8). (**D-E**) Surface expression of activation markers (CD25, CD44 and CD69), as measured by flow cytometry, on patient-derived CD4+ T cells, in autoimmune cohorts of (**D**) RA (n = 7) and (**E**) T1D (n = 8). (**F**) Experimental design of RA patient synovial fluid mononuclear cells (SFMCs). (**G**) IL-2, IL-10, IL-17, IFNγ and TNFα production by patient-derived SFMCs (n = 5). (**H**) Surface expression of activation markers (CD25, CD44 and CD69), as measured by flow cytometry, on patient-derived SFMCs (n = 5). All experiments were carried out using human samples. CD4+ T cells were activated with α-CD3 and α-CD28 for 24 h, in the presence and absence of ML-226, unless otherwise stated. Data are expressed as mean ± SEM.

To consolidate these findings, we assessed the efficacy of ABHD11 inhibition on synovial fluid mononuclear cells (SFMCs) isolated from the site of inflammation in a cohort of RA patients (Figure 4F). To this end, we observed targeted reductions in IL-17 and IFNγ production following ABHD11 inhibition, whilst IL-2, IL-10 and TNFα production remained unchanged (Figure 4G). In line with our earlier activation data (Figure 1D), CD4+ T cell activation remained intact following ABHD11 inhibition (Figure 4H), with a modest, but significant increase in cell size (Supplementary Figure 6E). Crucially, these changes did not result from compromised viability (Supplementary Figure 6F). Together, these findings demonstrate that ABHD11 inhibition retains its suppressive effect on T cell function in patients with autoimmune disease, including those present at the site of inflammation.

### ABHD11 inhibition delays the onset of murine type 1 diabetes

To support our findings in humans, we next investigated whether ABHD11 regulates T cell fate and function in a murine model of accelerated T1D. Here, we initially assessed the effect of ABHD11 inhibition on antigen-specific CD4+ T cells, whereby diabetogenic H2-Ag^7^-restricted BDC2.5 CD4+ T cells were stimulated *in vitro* in the presence of their cognate antigen^32^ (a hybrid insulin peptide [HIP]) and treated with the murine ABHD11 inhibitor, WWL222. There was a minimal reduction in proliferation observed following ABHD11 inhibition (Figure 5A). Despite a modest, but significant reduction in CD25 and CD69 expression on proliferating cells, the proportion of cells expressing these markers was unchanged (Figure 5B), again indicating that their activation is intact. Importantly, proinflammatory cytokine production is impaired by AHBD11 inhibition, with marked reductions in IFNγ, IL-2, IL-17 and TNFα (Figure 5C). Interestingly, this suppression appeared to be limited to proinflammatory cytokines, as we observed a striking increase in IL-10 production following ABHD11 inhibition (Figure 5C), which might indicate that ABHD11 inhibition induces an anti-inflammatory phenotype in antigen-specific T cells, rather than generally inhibiting effector function.

**Figure 5.**
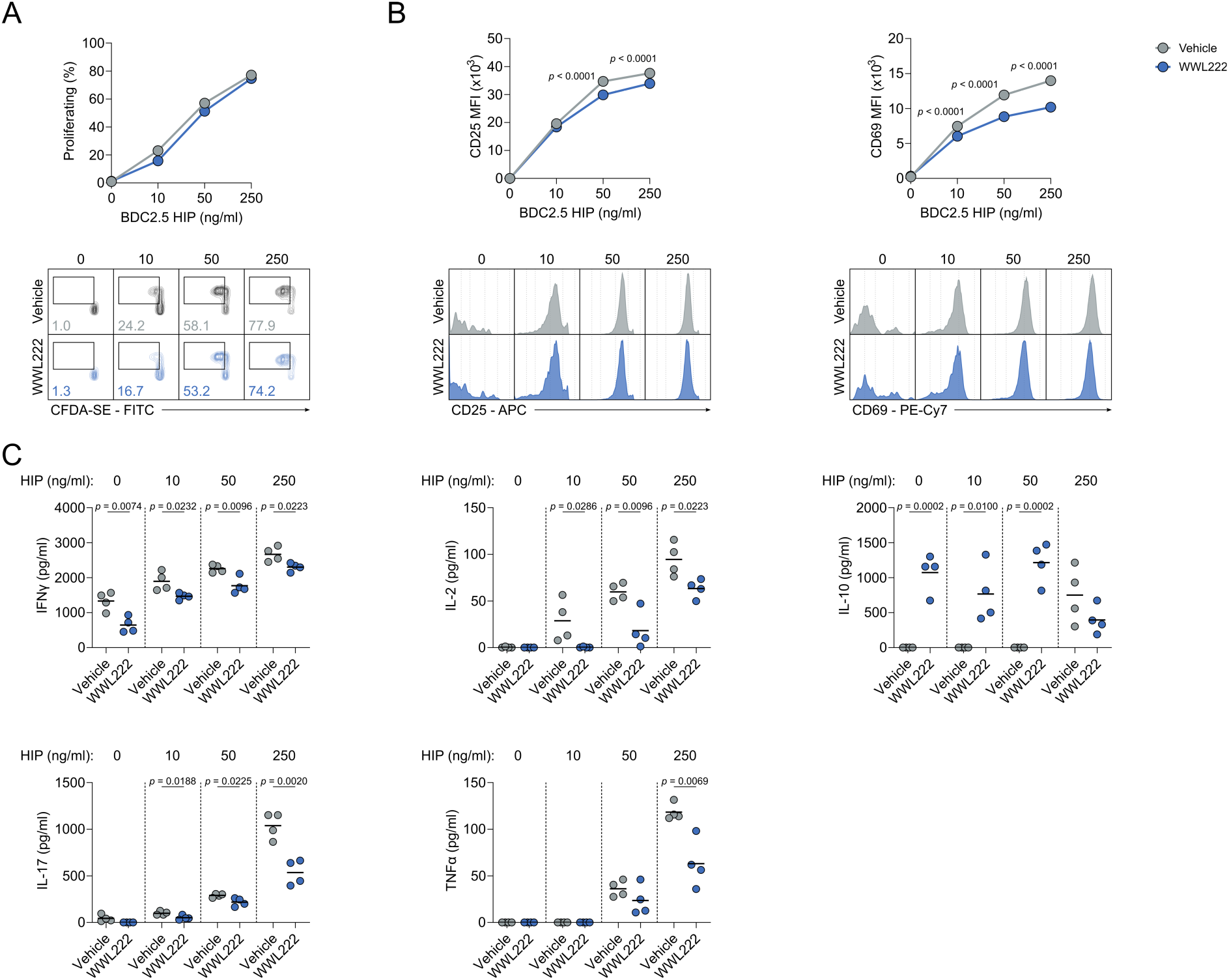
ABHD11 inhibition impairs the function of antigen-specific T cells. (**A**) Frequency of proliferating cells, as measured by flow cytometry using CFDA-SE, on antigen-specific T cells (n = 4). (**B**) Surface expression of activation markers (CD25 and CD69), as measured by flow cytometry, on antigen-specific T cells (n = 4). (**C**) IFNγ, IL-2, IL-10, IL-17 and TNFα production by antigen-specific T cells. All experiments were carried out using murine samples. BDC2.5 CD4+ T cells were activated with hybrid insulin peptides (HIPs), in the presence and absence of WWL222. Data are expressed as mean.

We furthered these investigations using an *in vivo* murine model of accelerated T1D. To this end, immunocompromised Rag-deficient mice were injected with 5×10^6^ peptide-activated BDC2.5 CD4+ T cells and WWL222 (or vehicle control), with the drug dose repeated daily for the course of the experiment (Figure 6A). Excitingly, ABHD11 inhibition by low-dose WWL222 treatment delayed the development of T1D (Figure 6B). At the end of the observation period, immune cells were harvested from the spleen of diabetic mice to elucidate the changes that underpin delayed onset of disease. Here, CD4+ T cells from WWL222-treated mice displayed impaired activation (Figure 6C), and produced significantly less IFNγ, TNFα and IL-2 (Figure 6D; Supplementary Figure 7A) versus those harvested from untreated mice. Additionally, TNFα was reduced in myeloid cell populations, meaning ABHD11 inhibition may skew the cytokine profile through multiple cell types (Supplementary Figure 7B-D). These data show that targeting ABHD11 can improve outcomes in the setting of autoimmunity, highlighting the significant potential of developing drugs that inhibit this metabolic node as novel treatment strategies in T cell-mediated autoimmunity.

**Figure 6.**
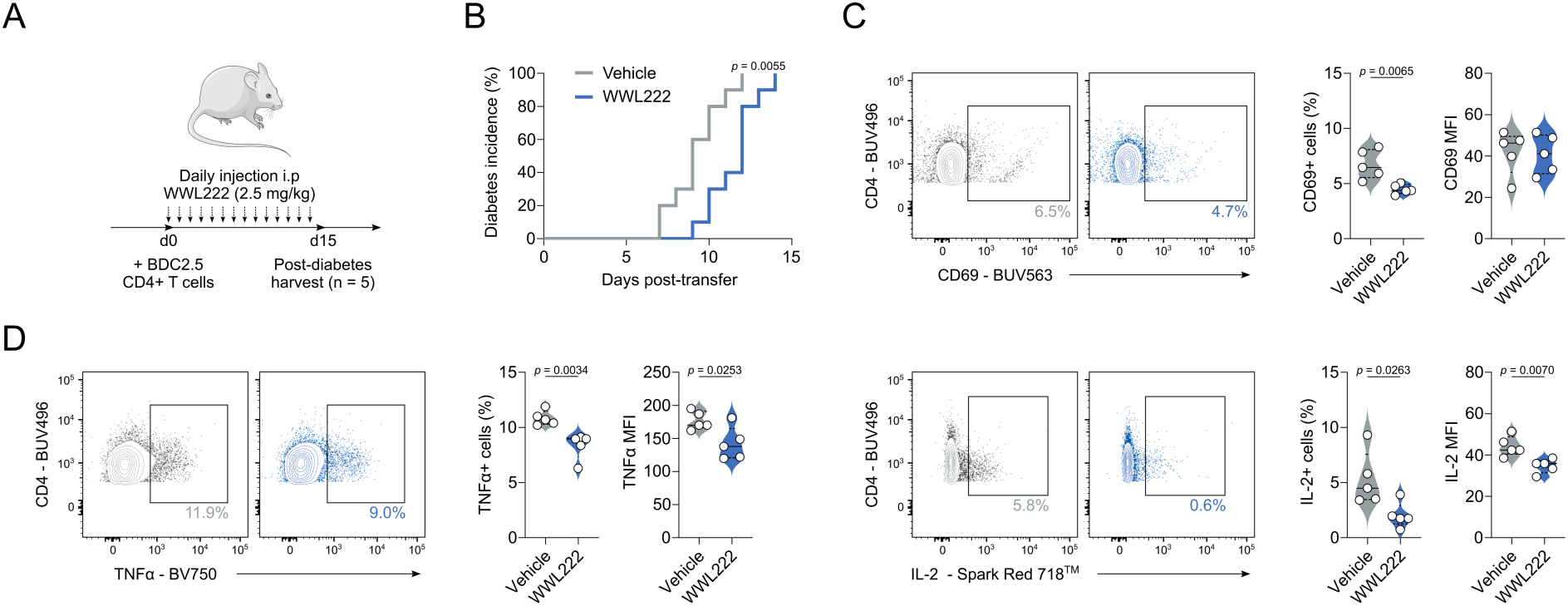
ABHD11 inhibition delays the onset of type 1 diabetes. (**A**) Schematic overview of *in vivo* diabetes adoptive transfer model using BDC2.5 HIP-activated BDC2.5 CD4+ T cells. (**B**) Diabetes incidence, as confirmed by blood glucose measurement > 13.9 mmol/L, in the presence and absence of daily injections i.p of 2.5 mg/kg WWL222. (**C**) Surface expression of CD69 as a measure of activation, as measured by flow cytometry, on CD4+ T cells (n = 5). (**D**) TNFα and IL-2 production, as measured by flow cytometry, by CD4+ T cells (n = 5). All experiments were carried out using murine samples. Mice were injected daily with the indicated dose of WWL222. Data are expressed as median ± interquartile range.

## Discussion

Reduced expression of ABHD11, a mitochondrial serine hydrolase, within patient CD4+ T cells is associated with clinical remission in RA^21^. However, beyond its role in maintaining the function of α-KGDH^22^, the function of ABHD11 in human T cells remains unknown, and as such, its significance in autoimmunity has yet to be explored. In this manuscript, we show that effector cytokine production is selectively impaired following ABHD11 inhibition, which is underpinned by the rewiring of mitochondrial metabolism. We thereby present ABHD11 as a potential drug target, through which to restrict autoreactive T cell function, for the treatment of T cell-mediated autoimmune disease.

A study from Bailey *et al*., first described that ABHD11 maintains functional lipoylation of the DLST subunit of α-KGDH^22^, which catalyses the conversion of KG to succinyl-CoA within the TCA cycle. Importantly, we demonstrate that α-KGDH activity is impaired in T cells treated with ML-226, a highly selective ABHD11 inhibitor, where there is a reduction in cellular succinate and a corresponding increase in the α-KG to succinate ratio. Accumulation of α-KG and subsequent formation of 2-HG can impair the activity of α-KG-dependent dioxygenases^33, 34, 35, 36^, which play an important role in epigenetic processes such as histone modifications, amongst other activities^37^. However, intracellular 2-HG levels were unchanged in T cells upon ABHD11 inhibition. Instead, the compromised TCA cycle drives the accumulation of lactate and acetyl-CoA. Despite evidence that high concentrations of acetyl-CoA enhance histone acetylation to subsequently shape T cell responses^38^, we did not observe any significant changes in histone acetylation following ABHD11 inhibition.

We instead reveal that ABHD11 inhibition restricts human T cell function through a SREBP2-mediated increase in oxysterol synthesis. SREBPs are considered to be the master regulators of lipid and sterol biosynthesis^39^. Although SREBP activity is required for the metabolic programming that prime T cells for growth and proliferation^40, 41^, the impact of augmented SREBP activation has yet to be fully explored in T cells. A recent study has described a lactate-SREBP2 signalling axis, whereby dendritic cells exposed to increased levels of lactate within the tumour microenvironment promoted cholesterol metabolism through a SREBP2-dependent pathway^28^. This appears to be driven by the concomitant change in extracellular pH, as an acidic pH (pH 6.8) has previously been shown to induce activation and nuclear translocation of SREBP2, leading to the expression of cholesterol biosynthesis genes^42^. Importantly, this relationship is not restricted to extracellular lactate levels, as the accumulation of lactate within cells has also been shown to promote SREBP2-mediated cholesterol biosynthesis^43^. In line with these studies, we show that following ABHD11 inhibition, there is an accumulation of intracellular lactate, with a concurrent upregulation of SREBP and sterol biosynthesis genes. Notably, this does not appear to be specific to the traditional mevalonate pathway in T cells, rather we show increased flux through a shunt pathway that produces 24,25-EC.

24,25-EC is one of the most efficacious of the oxysterols in the activation of LXR^29^, and we reveal that 24,25-EC levels are approximately 5-10x greater following AHBD11 inhibition. In keeping with a previous report that investigated the role of LXR activation in human CD4+ T cell function^31^, a significant proportion of the transcripts upregulated upon ABHD11 inhibition overlapped with those upregulated following exposure to a highly-selective LXR agonist GW3965. It is also important to note that LXR activation is closely integrated with SREBP signalling within inflammatory response programmes^44^. Typically, LXR activation is considered to be anti-inflammatory in a host of immune cell types^31, 45, 46^. Specifically, CD4+ T cells treated with LXR agonists display reduced proinflammatory function, including a reduction in their IL-17 production^31^. We show comparable reductions in T cell function following ABHD11 inhibition or treatment with a synthetic LXR agonist. 24,25-EC itself has also been shown to suppress immune cell function, limiting iNOS activation in LPS-activated monocytes in an LXR-dependent manner^47^. Thus heightened 24,25-EC levels, and downstream LXR activation, following ABHD11 inhibition culminate in suppressed effector function.

The relationship between metabolism and T cell effector function has been extensively explored during the past two decades^48, 49^. Indeed, aberrant metabolic programmes underpin the pathogenesis of several inflammatory conditions, including autoimmune disease, wherein unregulated T cell metabolism fuels a hyper-inflammatory phenotype^3, 4^. Given the relative ineffectiveness of current treatments^13, 50^, as well as the debilitating side effects that are often associated with these drugs^14^, novel approaches to resolve pathogenic tissue inflammation in autoimmune disease are highly attractive. Multiple pre-clinical studies have explored the prospect of modulating T cell metabolism to ameliorate inflammation and, thus, improve disease outcomes – for example, PKM2 tetramerisation^15^, suppressing OXPHOS using oligomycin^16^, and combined inhibition of glucose metabolism by 2-DG and metformin^17^. Despite these recent successes, the potential toxicity of targeting cellular metabolism at a systemic level remains an important consideration. In this manuscript, we targeted the typical immunometabolic profile of T cells by inhibiting ABHD11, with the aim of uncovering a druggable target that has wide applicability by overcoming the metabolic heterogeneity of autoimmune disease.

Crucially, ABHD11 inhibition retained its suppressive effect on T cell function in multiple settings of autoimmunity. Firstly, there was a comprehensive reduction in cytokine output when CD4+ T cells from RA or T1D patients were cultured *ex vivo* with ML-226. These findings were consolidated using a murine model of accelerated T1D, whereby ABHD11 inhibition suppressed antigen-specific T cell responses. Moreover, daily administration of WWL222 delayed the onset of diabetes, underpinned by a T cell-specific reduction in effector function, with only a minor effect on myeloid cell responses. Together, these findings encourage targeting ABHD11 to alleviate T cell-mediated autoimmune disease, thus providing the precedent for further pre-clinical studies.

In summary, we report that ABHD11 inhibition impairs T cell effector function. Our findings build upon a growing collection of studies that harness the importance of T cell metabolism for therapeutic benefit in autoimmunity. This work identifies a novel metabolic target to reshape conventional metabolic programmes and suppress the proinflammatory function of autoreactive T cells that are central to disease pathogenesis. Our manuscript supports the further development of ABHD11 inhibitors as a treatment for T cell-mediated autoimmunity.

## Methods

### Human samples

#### Healthy controls

Peripheral blood was collected from healthy, non-fasted individuals. Informed written consent and ethical approval was obtained from Swansea University Medical School Research Ethics Committee (SUMSRESC; 2022-0029). Peripheral blood mononuclear cells (PBMCs) were isolated via density gradient centrifugation using Lymphoprep™ (STEMCELL Technologies).

#### Autoimmune patient cohorts

PBMCs isolated from rheumatoid arthritis (RA; ethics RS18-055) and type 1 diabetes mellitus (T1D; ethics 12/WA/0033) patients were cryopreserved until use. SFMCs isolated from RA (ethics RS18-055) patients during arthroscopic knee surgery were cryopreserved until use. Cohort demographics can be found in Supplementary Tables 1-2.

### Human CD4+ T cell isolation and culture

Human CD4+ T cells (130-096-533) and CD4+ T effector (Teff) cells (130-094-125) were isolated by magnetic separation using the autoMACS^®^ Pro Separator as per the manufacturer’s instructions (Miltenyi). T cells were activated with plate-bound anti-CD3 (2 μg/ml; OKT3; BioLegend) and soluble anti-CD28 (20 μg/ml; CD28.2; BioLegend) in human plasma-like medium (HPLM; Gibco) at 37°C in 5% CO_2_-in-air for 24 h. To prevent impaired T cell activation, culture media was supplemented with 10% dialysed fetal bovine serum (FBS; Fisher Scientific) after 3 h. T cells were treated with ML-226 (10 μM; Cambridge Bioscience), unless otherwise stated.

#### Phenocopy assays

For LXR activation experiments, T cells were activated in the presence and absence of GW3965 (2 μM; Merck). For lactate phenocopy experiments, T cells were activated in the presence and absence of lactic acid (10 mM; Merck).

### Murine CD4+ T cell isolation and culture

#### Mice

BDC2.5 TCR transgenic NOD mice (NOD.Cg-Tg(TcraBDC2.5,TcrbBDC2.5)1Doi/DoiJ; 004460) and Rag1-/-NOD mice (NOD.129S7(B6)-Rag1tm1Mom/J; 003729) were both purchased from Jackson Laboratory^51, 52, 53^. All mice received water and irradiated food ad libitum and were housed at Cardiff University in specific-pathogen-free Scantainers with 12 h light–dark cycles. All animal experiments were approved by the Cardiff University ethical review process and conducted under UK Home Office licence in accordance with the UK Animals (Scientific Procedures) Act 1986 and associated guidelines.

#### In vitro T cell differentiation assay

Splenic CD4+ T cells were isolated from C57BL/6 mice (aged 8-12 weeks) by MACS magnetic separation (Miltenyi) and cultured in RPMI-1640 (Gibco) media at 37°C in 5% CO_2_-in-air for 4 d. For Th17 polarising conditions, Iscove’s Modified Dulbecco’s Medium (IMDM; Gibco) was used for optimal Th17 differentiation. CD4+ T cells were activated with plate-bound anti-CD3 (1 μg/ml; 145-2C11; R&D Systems) and soluble anti-CD28 (10 μg/ml; 37.51; eBioscience). Polarisation conditions were as follows: Th1 (IL-12 [20 ng/ml]), Th2 (IL-4 [40 ng/ml]), Th17 (TGFβ [1 ng/ml], IL-6 [20 ng/ml], IL-23 [20 ng/ml], anti-IL-2 [10 µg/ml, JES6–1A12]). All cytokines were purchased from R&D Systems. T cells were treated with WWL-222 (10 µM) or DMSO vehicle control.

#### In vitro antigen presentation assay

Splenic CD4+ T cells were isolated from BDC2.5 NOD mice as per the manufacturer’s instructions (MojoSort™ Mouse CD4 T cell Isolation Kit; BioLegend) and subsequently labelled with 0.5 μM CFDA-SE (Vybrant CFDA-SE Cell Tracer Kit; Invitrogen). Splenic antigen-presenting cells (APCs) were depleted of T cells through incubating cells with anti-Thy1 antibody (50 μg/ml, M5/49.4.1; BioXcell) and 1:20 rabbit complement (Merck) over 1 h at 37°C. 1×10^5^ BDC2.5 CFDA-SE-labelled CD4+ T cells were co-cultured 1:1 with APCs in the presence of BDC2.5 hybrid insulin peptide (DLQTLALWSRMD)^32^ for 48 h prior to harvest. Proliferating BDC2.5 CD4+ T cells (CFDA-SE^low^) were analysed by flow cytometry.

#### In vivo diabetes adoptive transfer

BDC2.5 CD4+ T cells were activated and isolated as previously described. 5×10^6^ BDC2.5 CD4+ T cells were i.v. adoptively transferred into 3-4 week-old Rag1-/-NOD recipients. Recipient mice were injected with 2.5mg/kg of WWL222 daily i.p. from the day of T cell transfer until termination. Mice were screened for glycosuria daily (Diastix; Bayer) with diabetes confirmed by blood glucose measurement (> 13.9 mmol/L).

### Flow cytometry

#### Gating strategy

Flow cytometry was performed on T cells following cell culture. Cell death was monitored using DRAQ7™ (1μM, DR71000; Biostatus), unless otherwise stated, and dead cells were excluded from analysis. Cell doublets were also excluded from analysis. A representative gating strategy can be found as Supplementary Figure 8.

#### Surface staining

For human T cells, surface staining was performed at room temperature (RT) for 15 min in the dark. Antibodies were used as follows: anti-CD25 (Pacific Blue, mIgG1κ, BC96, 302627), anti-CD25 (PE-Vio 615, rhIgG1, REA570, 130-123-035; Miltenyi), anti-CD44 (FITC, rhIgG1, REA690, 130-113-341; Miltenyi), anti-CD44 (Pacific Blue, mIgG1κ, BJ18, 338823), anti-CD69 (APC, mIgG1κ, FN50, 310910), anti-CD69 (FITC, mIgG1κ, FN50, 310904). Antibodies were purchased from BioLegend, unless otherwise stated.

For murine splenic T cells from C57BL/6 mice, cell death was monitored using the Zombie Aqua Fixable Viability Kit (BD).

For murine splenic T cells from BDC2.5 TCR transgenic NOD mice, single cell suspensions were incubated with TruStain FcX™ Fc Receptor Blocking Solution (clone 93; BioLegend) for 10 min at 4°C prior to staining for surface markers for 30 min at 4°C. Antibodies for were used as follows: CD4 (BUV496, LewIgG2b, GK1.5, 612952; BD), CD8 (PerCP/Cy5.5, rIgG2aκ, 53-6.7, 100734), CD11b (Brilliant Violet 510™, rIgG2bκ, M1/70, 101263), CD11c (BUV661, ahIgG2, N418, 750449; BD), CD19 (APC-Cy7, rIgG2aκ, 1D3, 152412), CD25 (APC, rIgG1, PC61, 102012), CD40 (BUV615, LouIgG2aκ, 3/23, 751646; BD), CD44 (BUV805, rIgG2bκ, IM7, 741921; BD), CD62L (BUV395, rIgG2aκ, MEL-14, 569400; BD), CD69 (BUV563, ahIgG1λ3, H1.2F3, 612952; BD), CD80 (AlexaFluor® 594, rrIgG, 2740B, FAB7401T-100ug; Bio-Techne), CD86 (BUV737, rIgG2bκ, PO3, 741757; BD), MHC-I-[H-2kd] (Brilliant Violet 421™, mIgG2aκ, SF1-1.1, 116623) and MHC-II-[I-AK] (PE, mIgG2aκ, 10-3.6, 109908). Antibodies were purchased from BioLegend, unless otherwise stated. Cell death was monitored using either the Zombie Aqua Fixable Viability Kit or Fixable Viability Stain 575V (both BD).

#### Intracellular staining

For human T cells, intracellular markers were stained using the eBioscience™ Foxp3 / Transcription Factor Staining Buffer Set (00-5523-00) and cell death monitored using the eBioscience™ Fixable Viability Dye eFluor™ 506 (65-0866-14) as per the manufacturer’s instructions (both Invitrogen). 4 h prior to intracellular staining, cells were activated with PMA (50 ng/ml; Merck) and ionomycin (500 ng/ml; Merck), and protein transport blocked using eBioscience Protein Transport Inhibitor Cocktail (Invitrogen). Following surface staining, cells were fixed for 30 min at RT before staining overnight at 4°C in permeablisation buffer. Antibodies used were purchased from BioLegend, unless otherwise stated: anti-IFNγ (FITC, mIgG1κ, 4S.B3, 502506), anti-TNFα (PE, mIgG1κ, Mab11, 502909).

For murine T cells from C57BL/6 mice spleens, intracellular markers were stained using the Cytofix/Cytoperm Fixation/Permeabilisation kit (BD) as per the manufacturer’s instructions. 4 h prior to intracellular staining, cells were activated with PMA (50 ng/ml) and ionomycin (500 ng/ml), and protein transport blocked using monensin (3 μM; all Merck). Following viability staining, cells were fixed and permeabilised and stained with CD4 (Brilliant Violet 758™, rIgG2aκ, RM4-5, 100551; BioLegend), IL-13 (FITC, rIgG1κ, eBio13A, 53-7133-82; Invitrogen), IL-17A (PE, rIgG1κ, TC11-18H10, 561020; BD Pharmigen) and IFNγ (eFluor™ 450, rIgG1κ, XMG1.2, 48-7311-82; Invitrogen).

For murine T cells from BDC2.5 TCR transgenic NOD mice spleens, intracellular markers were stained using the Cytofix/Cytoperm Fixation/Permeabilisation kit (BD) as per the manufacturer’s instructions. 3 h prior to intracellular staining, cells were activated with PMA (50 ng/ml) and ionomycin (500 ng/ml), and protein transport blocked using GolgiPlug (BD). Following surface staining, cells were fixed for 20 min at RT before permeabilisation. Cells were incubated with TruStain FcX™ as previously described prior to staining for intracellular cytokines. Antibodies used were purchased from BioLegend, unless otherwise stated: IL-2 (Spark Red™ 718, rIgG2bκ, JES6-5H4, 503852), IL-6 (APC, rIgG1κ, MP5-20F3, 503852), IL-10 (Brilliant Violet 605™, rIgG2bκ, JES5-16E3, 505031), IL-12/23 (PE-Cy7, rIgG2κ, C15.6, 505210), IL-17a (Brilliant Violet 786™, rIgG2κ, TC11-18H10.1, 506928), IFNγ (Brilliant Violet 650™, rIgG1κ, XMG1.2, 505832) and TNFα (Brilliant Violet 750™, rIgG1κ, MP6-XT22, 506358).

#### Puromycin incorporation

Protein translation was assessed using anti-puromycin (AlexaFluor® 488, 12D10, MABE343-AF488; Merck). Puromycin (10 μM; Merck) was added 15 min prior to the end of 4 h and 24 h T cell activation. Cells were washed in ice-cold PBS before intracellular staining was performed using Inside Stain Kit (Miltenyi) as per the manufacturer’s instructions. Cells were fixed for 20 min at RT, permeabilised for 15 min at RT, before staining for 1 h at 4°C in permeabilisation buffer.

#### Mitochondrial characteristics

For mitochondria content and membrane potential, cells were incubated with MitoTracker™ Green FM (100 nM; M7514, ThermoFisher) or TMRE (50 nM; ab113852; Abcam) for 20 min at 37°C, respectively. For mitochondrial ROS staining, cells were incubated with MitoSOX™ Red (5 μM; M36008, ThermoFisher) for 20 min at 37°C.

#### Purity

Human CD4+ T cell purity was monitored using anti-CD3 (Brilliant Violet 570™, mIgG1κ, UCHT1, 300436) and anti-CD4 (AlexaFluor® 647, mIgG2b, OKT4, 317422, BioLegend). CD4+ effector T cell purity was monitored using anti-CD4 (AlexaFluor® 647, mIgG2b, OKT4, 317422, BioLegend), anti-CD45RA (Brilliant Violet 605™, mIgG2b, HI100, 304134, BioLegend), anti-CD45RO (FITC, mIgG2a, UCHL1, 304204, BioLegend) and anti-CD197 (Pacific Blue, mIgG2a, G043H7, 353210, BioLegend). Percentage purity was consistently > 90%.

#### Analysis

Human T cells were acquired on a Novocyte 3000 (Agilent). Murine T cells were acquired on a Novocyte 3000 (Agilent), FACS Canto II or a Symphony A3 Cell Analyser (both BD). Analysis was performed using FlowJo version 10 (TreeStar).

### Enzyme linked immunosorbent assay

Cell-free supernatants were analysed for human IL-2 (DY202), IL-10 (DY217B), IL-17 (DY317), IFNγ (DY285B) and TNFα (DY210) by DuoSet ELISA as per the manufacturer’s instructions (R&D Systems). 96-well half-area plates were coated with capture antibody overnight at 4°C. Cell-free supernatants were diluted to an appropriate concentration and incubated for 2 h at RT with gentle agitation, followed by 2 h with the kit-specific detection antibody, and finally 20 min with streptavidin-horse radish peroxidase. The plate was then incubated with a 1:1 mixture of hydrogen peroxide and tetramethylbenzoic acid (555214; BD Biosciences) at RT. Absorbance was measured at 450 nm following the addition of sulfuric acid (Merck) to each well. All values were corrected to the blank.

### Immunoblot

T cells were lysed in PhosphoSafe Extraction Buffer (Merck). Cell lysate proteins were quantified, denatured and separated using SDS-PAGE. Polyvinylidene difluoride membranes were probed with antibodies targeting ABHD11 (PA5-54962) and acetylated lysine (MA5-33031). All antibodies were purchased from Invitrogen and used at a 1:1000 dilution. Protein loading was monitored using β-actin (ab8226, Abcam).

### Gene expression analysis

#### RNA preparation

RNA was extracted using RNeasy® Mini Kit columns (74014; Qiagen) as per the manufacturer’s guidelines. RNA purity was assessed using a Nanodrop™ spectrophotometer and measured A260/280 and A260/230 ratios were typically between 1.8 – 2.2.

#### qPCR

cDNA was prepared from 500 ng of RNA using the High-Capacity cDNA Reverse Transcription Kit (4368813; ThermoFisher) as per the manufacturer’s instructions. qPCR reactions were prepared using Fast SYBR® Green Master Mix (4385612; ThermoFisher) as per the manufacturer’s guidelines. All gene expression analyses were normalised to *RPL19*. Primer sequences can be found in Supplementary Table 3.

### Metabolic analysis

Metabolic analysis was performed using the Seahorse Extracellular Flux Analyzer XFe96 (Agilent) following cell culture. T cells were resuspended in RPMI phenol red free media supplemented with glucose (10 mM), glutamine (2 mM) and pyruvate (1 mM; all Agilent). T cells were seeded onto a Cell-Tak (354240; Corning) coated microplate allowing the adhesion of T cells. Mitochondrial and glycolytic respiratory parameters were measured using OCR (pmoles/min) and ECAR (mpH/min), respectively. Injections included: oligomycin (1 μM), FCCP (1 μM), rotenone (1 μM) and antimycin A (1 μM) and monensin (20 μM). All chemicals were purchased from Merck, unless otherwise stated.

### Liquid-chromatography mass-spectrometry (LC-MS)

T cells were washed twice with ice-cold PBS and lysed in methanol, acetonitrile and water (v/v 5:3:2) following cell culture. Chromatographic separation of metabolite extracts was done using a ZIC-pHILIC column (SeQuant; 150 mm × 2.1 mm, 5 µm; Merck) and ZIC-pHILIC guard column (SeQuant; 20 mm × 2.1 mm; Merck) coupled to Vanquish HPLC system (ThermoFisher). A gradient programme was employed, using 20 mM ammonium carbonate (pH 9.2, 0.1 % v/v ammonia, 5 µM InfinityLab deactivator (Agilent)) as mobile phase A and 100% acetonitrile as mobile phase B. Elution started at 20% A (2 min), followed by a linear increase to 80% A for 15 min and a final re-equilibration step to 20% A. Column oven was set to 45 °C and flow rate to 200 µl min^−1^.

Metabolite profiling and identification was achieved using a Q Exactive Plus Orbitrap mass spectrometer (ThermoFisher) equipped with electrospray ionization. Polarity switching mode was used with a resolution (RES) of 70,000 at 200 m/z to enable both positive and negative ions to be detected across a mass range of 75 to 1,000 m/z (automatic gain control (AGC) target of 1 × 10^6^ and maximal injection time (IT) of 250 ms).

Data analysis was undertaken in Skyline (version 23.1.0.455)^54^. Identification was accomplished by matching accurate mass and retention time of observed peaks to an in-house library generated using metabolite standards (mass tolerance of 5 ppm and retention time tolerance of 0.5 min).

### Stable isotope tracer analysis (SITA) by LC-MS

Isolated human CD4+ T cells were incubated with universally heavy-labelled ^13^C_6_-glucose (11.1 mM; Cambridge Isotopes) in glucose-free RPMI (Gibco), or ^13^C_5_-glutamine (2 mM; Cambridge Isotopes) in glutamine-free RPMI (Gibco), and activated in the presence and absence of ML-226, as previously described. T cells were then washed twice with ice-cold PBS and lysed in methanol, acetonitrile and water (v/v 5:3:2). Metabolite extraction and subsequent LC-MS analysis were performed as previously described. For tracing analysis, integration of each isotopologue was manually verified.

### Sterol analysis

T cell pellets (> 4×10^6^ cells) were snap-frozen in liquid nitrogen. Sterol extraction, hydrolysis, isolation and analysis was performed according to previously described methods^55^.

### RNA-Seq

#### Sample preparation

T cell pellets were snap-frozen in liquid nitrogen. RNA was extracted using RNeasy® Mini Kit columns (74014; Qiagen) as per the manufacturer’s guidelines. For each sample, 2 μg of total RNA was then used in Illumina’s TruSeq Stranded mRNA Library kit (#20020594). Libraries were sequenced on Illumina NovaSeq 6000 as paired-end 150-nt reads.

#### Analysis

Fastq files were quality assessed and trimmed using FastP(v0.23.1)^56^, before reads were mapped to the genome GRCh38 using STAR (Spliced Transcripts Alignment to a Reference; v2.7.9a)^57^ with 2-pass method and multimapping set to 1. Featurecounts(v2.0.3)^58^ was used to generate count files for each sample, with counting performed at the gene level. Differential gene expression analysis was calculated via eBayesian fit of TMM (Trimmed Mean of M-values) using an EdgeR workflow^59^ of Limma-Voom(v3.58.1)^60^.

Genes were filtered for an adjusted p-value < 0.05 and over representation analysis was performed using gProfiler (v.e111.eg58.p18.f463989d)^61^ for Gene Ontology terms for Biological Processes (GO:BP). ReViGo(v1.8.1)^62^ was used to reduce the repetition of canonical pathway terms. GeneSet enrichment analysis was performed using genekitR (v1.2.5)^63^. Protein-protein interaction was assessed using StringDB(v12.0)^64^, whereby differentially expressed genes were treated as if they were fully transcribed into proteins. Transcription factor enrichment analysis was performed using X2Kweb (v14/22 Appyter)^65^, which infers upstream regulatory networks from the differentially-expressed gene signature. For all enrichment analyses, differentially-expressed genes were separated into up- and downregulated sets.

### α-ketoglutarate dehydrogenase activity assay

T cell α-ketoglutarate dehydrogenase activity was measured following culture using the α-Ketoglutarate Dehydrogenase Activity Colorimetric Assay Kit (MAK189; Merck) as per the manufacturer’s instructions. T cells were lysed with the α-KGDH Assay Buffer provided, before commencing the reaction with α-KGDH developer and α-KGDH substrate. Absorbance was measured at 450 nm until the absorbance value for the most active sample exceeded that of the highest NADH standard concentration. All values were corrected to the blank.

### Lactate assay

Extracellular L-lactate concentrations were measured using L-Lactate Assay Kit I (Eton Bioscience, USA) as per the manufacturer’s instructions. Cell-free supernatants were diluted to an appropriate concentration prior to addition of L-Lactate Assay Solution and incubation at 37°C for 30 min. Absorbance was measured at 490 nm after the addition of acetic acid (0.5 M; Merck) to each well. All values were corrected to the blank.

### Statistical analysis

Statistical analysis was performed using GraphPad Prism version 10 (USA). Data are represented as the mean ± or + standard error of the mean (SEM). The one-sample Kolmogorov-Smirnoff test was used to test normality. Where no substantial deviations from normality were observed, it was considered appropriate to use parametric statistics. All experiments have replicate sample sizes of ≥ n = 3 and significant values were taken as * *p* ≤ 0.05, ** *p* ≤ 0.01, *** *p* ≤ 0.001 and **** *p* < 0.0001.

### Data availability

All data are available upon request and can be found within the manuscript and supplementary information.

## Acknowledgements

We thank W. J. Griffiths for useful discussion, and all blood donors for their contribution to this work. We would also like to acknowledge Active Motif for their support. Y.R.J. is funded by a Swansea University Research Excellence Scholarship. A.H.U., F.B. and D.S. are funded by Cancer Research UK core funding to the CRUK Scotland Institute. L.C.D. is funded by an MRC New Investigator Research Grant (MR/Y013816/1). J.A.N is supported by a Wellcome Senior Clinical Research Fellowship (215477/Z/19/Z). JVV is supported by a Cancer Research UK - Career Development Fellowship [RCCCDF-Nov23/100001] and by a Lord Kelvin/Adam Smith (LKAS) Leadership Fellowship from the University of Glasgow. J.G.M. is funded by National Institutes for Health (NIH) research grants (UL1TR003163, 1P30DK127984 and P01HL160487). G.W.J. is funded by a Versus Arthritis Career Development Fellowship (20305). J.A.P is supported by an MRC Career Development Award (MR/T010525/1). E.E.V is supported by a Diabetes UK RD Lawrence Fellowship (17/0005587) and by Cancer Research UK (C18281/A29019). This work was funded by an MRC New Investigator Research Grant (MR/X000095/1) awarded to N.J.

## Authorship Contribution

B.J.J., Y.R.J., F.M.P-G., C.M., M.D.H., A.H.U., F.B., S.E., A.B., J.D., J.B., G.D.V., J.A.P., and N.J. performed the experiments. B.J.J, Y.R.J, D.K.F., L.V.S., A.E.H., D.S., J.V.V., G.W.J., J.A.P., E.E.V., and N.J. designed the experiments. B.J.J., Y.R.J., I.A.P., C.M., M.D.H., D.S., A.H.U., S.E., J.V.V., G.D.V., J.A.P., and N.J. analysed the data. A.H., D.J.V., U.F., and J.A.P. provided access to clinical samples. B.J.J., Y.R.J., J.G.C., J.B., L.C.D., M.N., D.K.F., L.V.S., B.F.C., A.E.H., J.A.N., U.F., D.S., J.V.V., J.G.M., G.W.J., J.A.P., E.E.V. and N.J. provided intellectual discussion. B.J.J., Y.R.J, J.A.P, E.E.V. and N.J. wrote the manuscript. All authors critically revised and approved the manuscript.

---

**Supplementary Figure 1.**
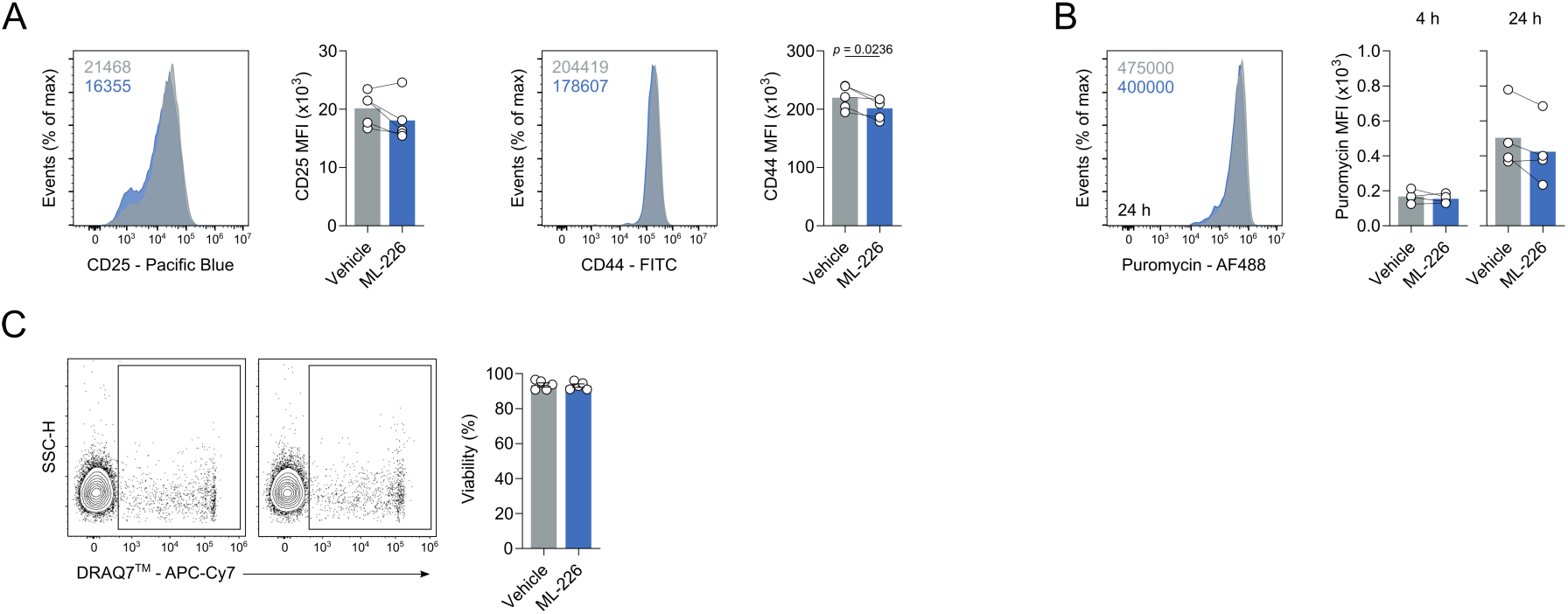
AHBD11 inhibition does not significantly alter T cell size, protein translation and viability. (**A**) Surface expression of activation markers (CD25 and CD44), as measured by flow cytometry, on CD4+ effector T cells (n = 5). (**B**) Puromycin incorporation, as measured by flow cytometry, in CD4+ effector T cells (n = 4). (**C**) Cell viability, as determined by DRAQ7^®^, in CD4+ effector T cells (n = 5). All experiments were carried out using human samples. CD4+ T cells were activated with α-CD3 and α-CD28 for 24 h, in the presence and absence of ML-226, unless otherwise stated. Data are expressed as either: mean, with paired dots representing biological replicates; or mean ± SEM.

**Supplementary Figure 2.**
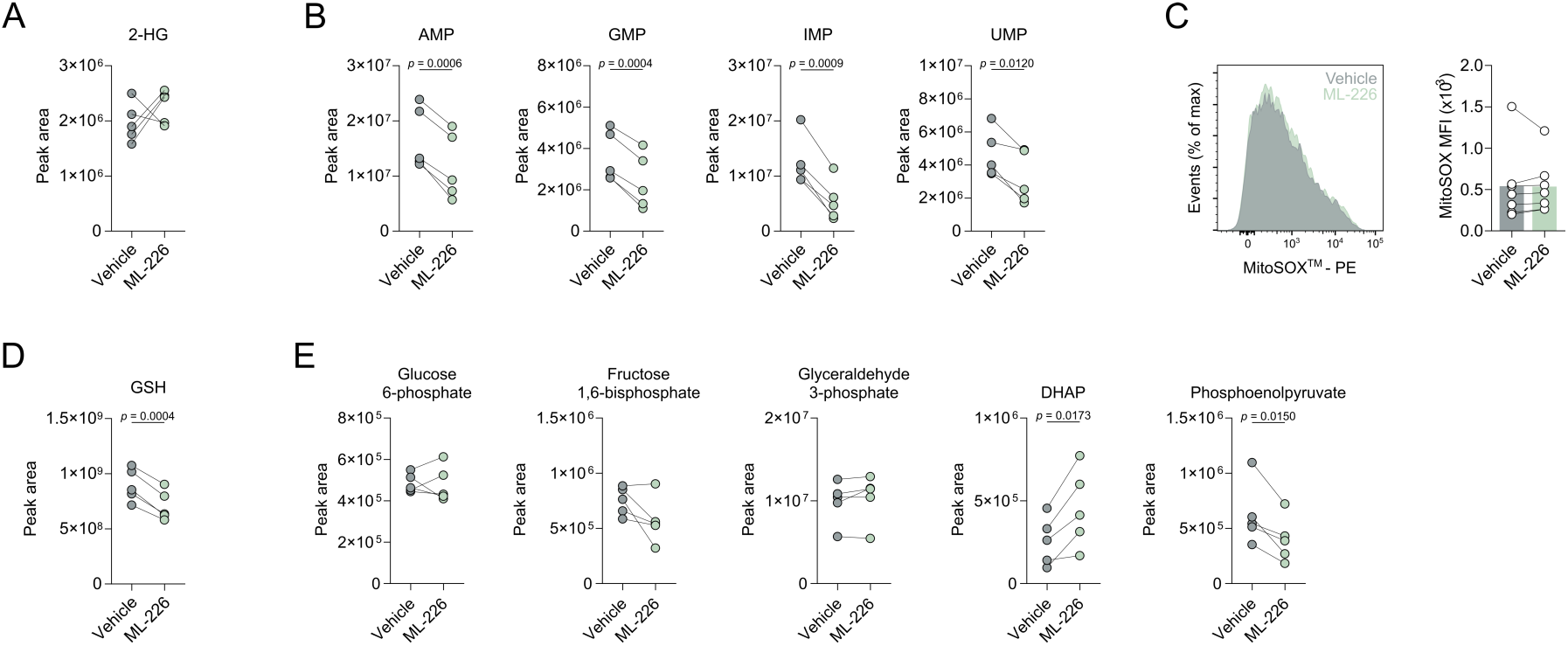
ABHD11 inhibition reduces intracellular monophosphate nucleotides. (**A**) Intracellular levels of 2-hydroxyglutarate (2-HG) in CD4+ T cells (n = 5). (**B**) Intracellular levels of selected monophosphate nucleotides in CD4+ effector T cells activated with α-CD3 and α-CD28 for 24 h, in the presence and absence of ML-226 (n = 5). Metabolites include: inosine monophosphate, adenosine monophosphate, guanosine monophosphate and uridine monophosphate. (**C**) Mitochondrial ROS levels, as determined by MitoSOX™ Red, in CD4+ effector T cells (n = 7). (**D**) Intracellular levels of glutathione in CD4+ effector T cells (n = 5). (**E**) Intracellular levels of selected glycolytic intermediates in CD4+ effector T cells (n = 5). Metabolites include: glucose 6-phosphate, fructose 1,6-bisphosphate, glyceraldehyde 3-phosphate, dihydroxyacetone phosphate (DHAP) and phosphoenolpyruvate. All experiments were carried out using human samples. CD4+ T cells were activated with α-CD3 and α-CD28 for 24 h, in the presence and absence of ML-226, unless otherwise stated. Data are expressed as mean, with paired dots representing biological replicates.

**Supplementary Figure 3.**
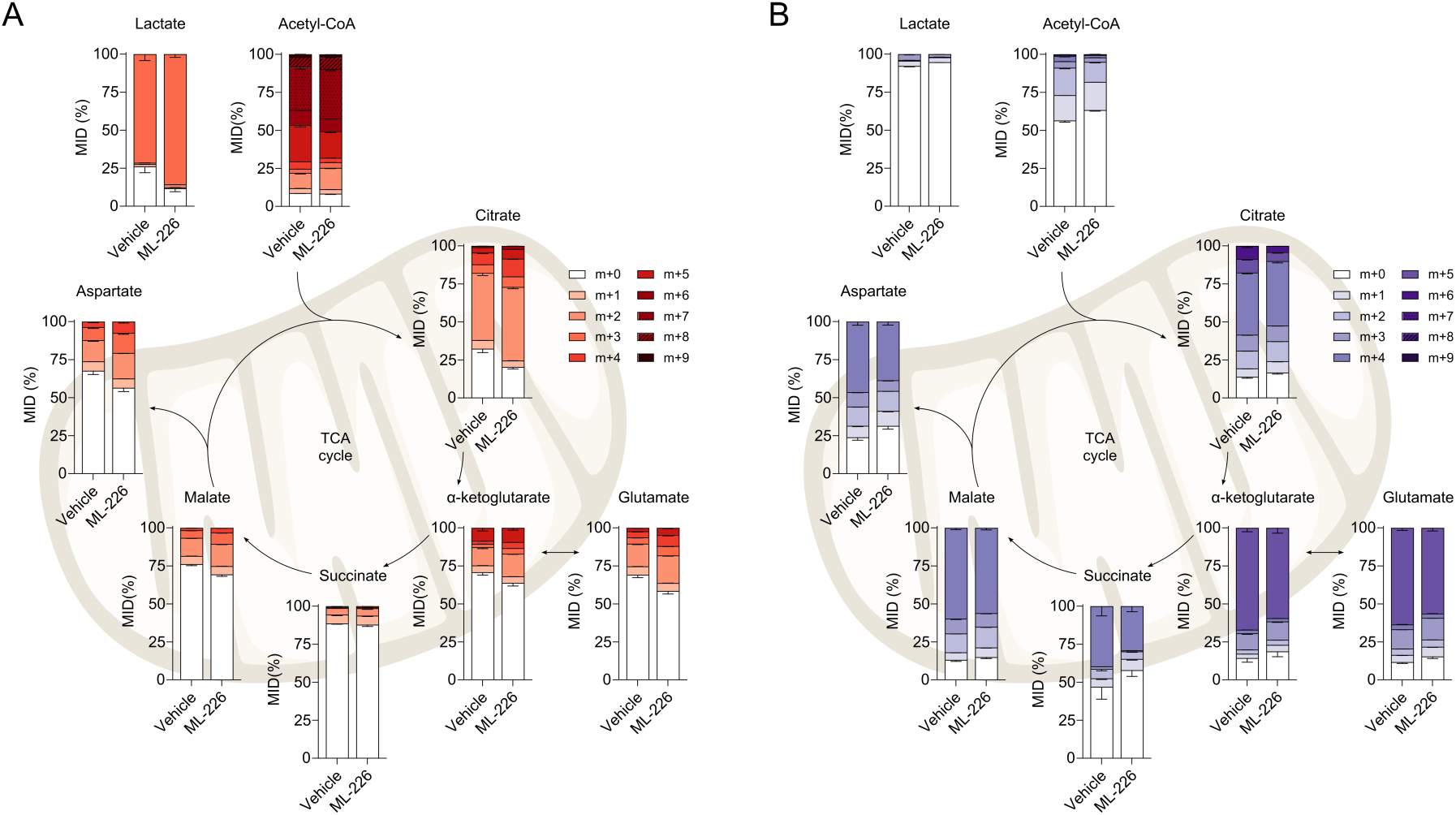
ABHD11 inhibition alters glucose and glutamine utilisation. (**A**) Stable isotope tracing of uniformly labelled ^13^C_6_-glucose into the TCA cycle and related intermediates in CD4+ effector T cells (n = 6). (**B**) Stable isotope tracing of uniformly labelled ^13^C_5_-glutamine into the TCA cycle and related intermediates in CD4+ effector T cells (n = 6). Metabolites include: lactate, acetyl-CoA, citrate, α-ketoglutarate, glutamate, succinate, malate and aspartate. Mass isotopologue distribution (MID) represented as the proportion of the metabolite pool. All experiments were carried out using human samples. CD4+ T cells were activated with α-CD3 and α-CD28 for 24 h, in the presence and absence of ML-226, unless otherwise stated. Data are expressed as mean ± SEM.

**Supplementary Figure 4.**
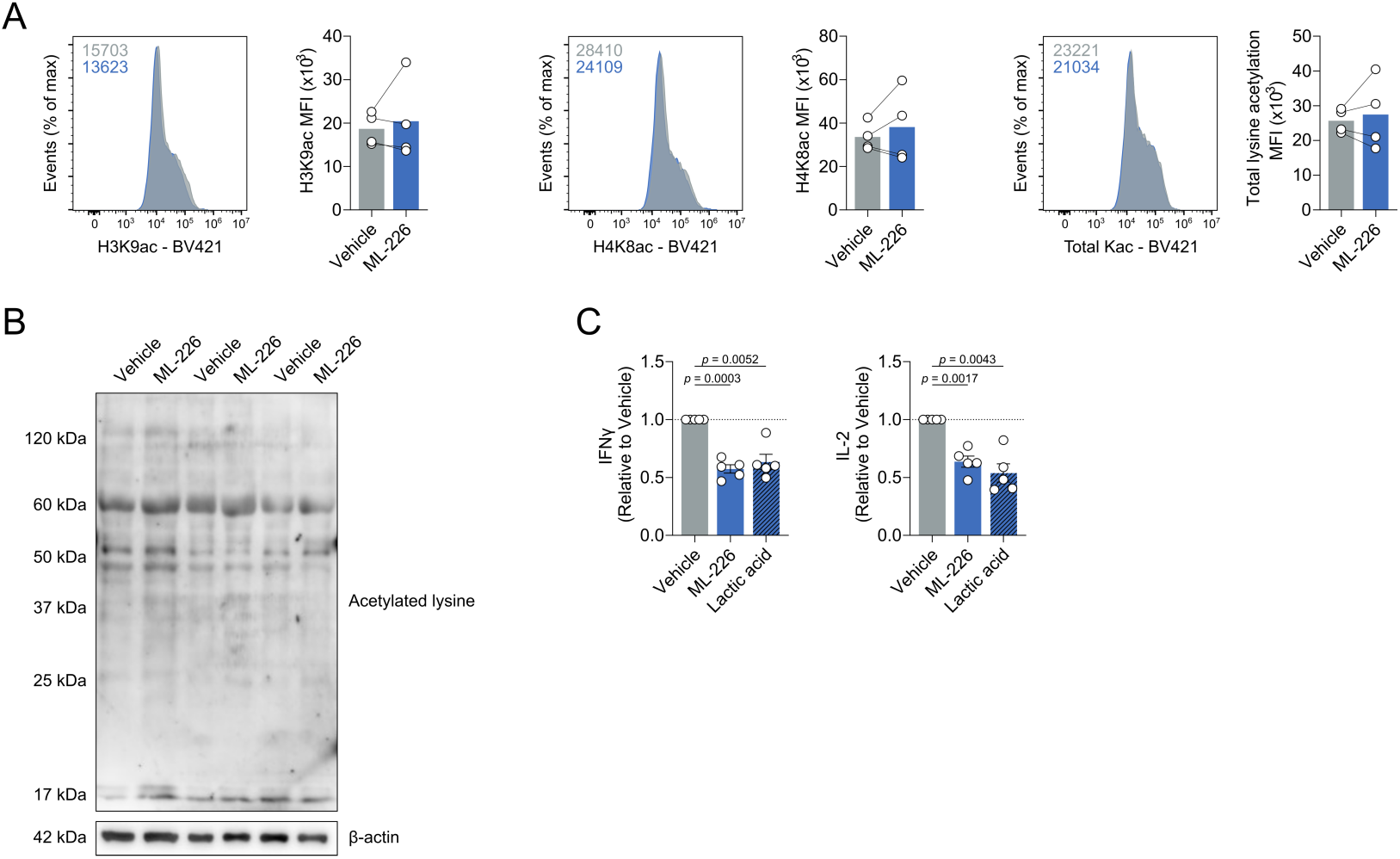
ABHD11 inhibition has contrasting effects on epigenetic modifications. (**A**) Intracellular histone acetylation levels, as measured by flow cytometry, in CD4+ effector T cells (n = 4). Histone acetylation measured on: H3K9 and H4K8. (**B**) Total lysine acetylation in CD4+ T cells (n = 3). Protein loading assessed using β-actin. (**C**) IL-2 and IFNγ production by CD4+ T cells, activated in the presence and absence of ML-226 or lactic acid (n = 5). All experiments were carried out using human samples. CD4+ T cells were activated with α-CD3 and α-CD28 for 24 h. Data are expressed as either: mean, with paired dots representing biological replicates; or mean ± SEM.

**Supplementary Figure 5.**
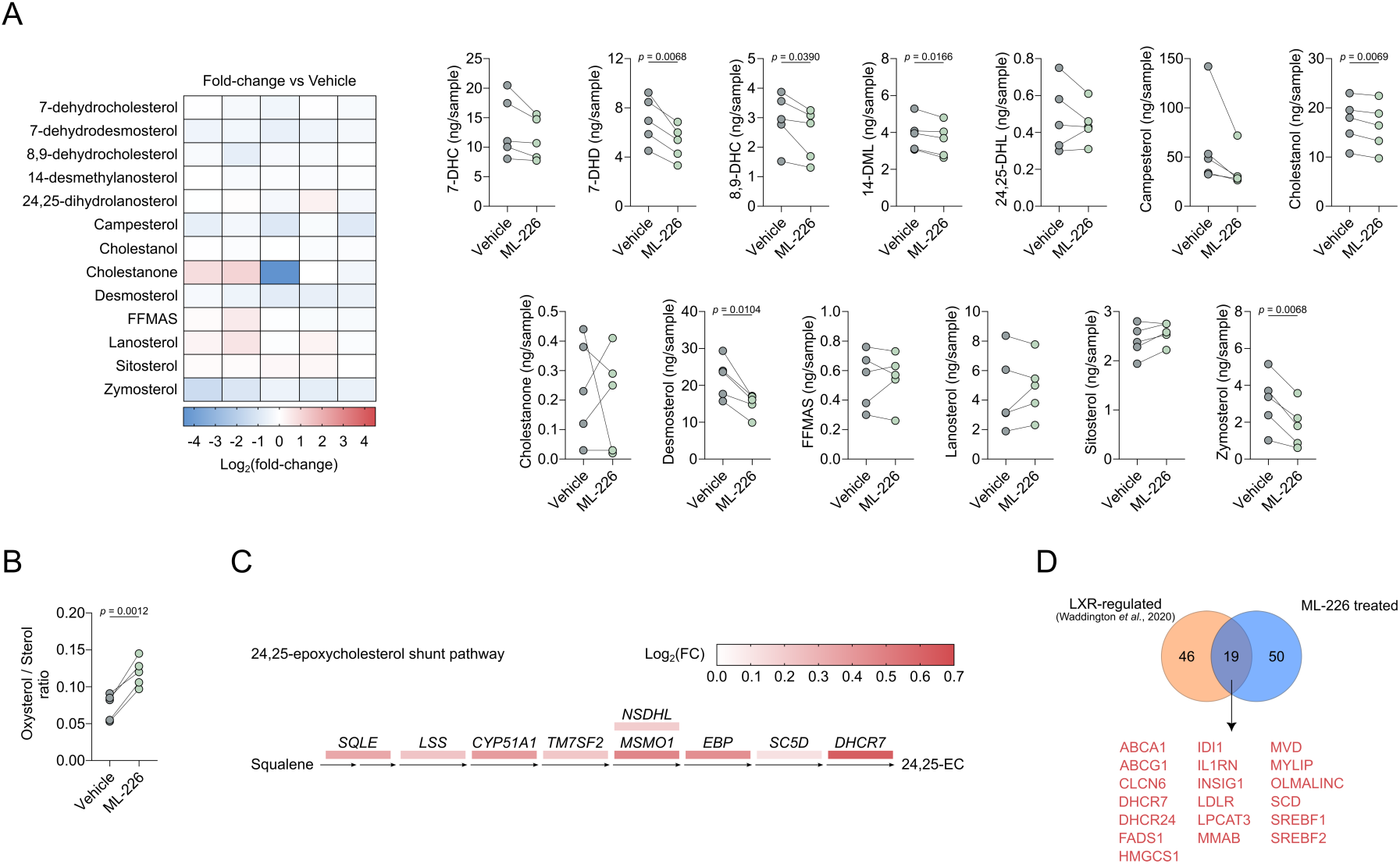
ABHD11 inhibition reduces non-oxygenated sterol levels and drives activation of a mevalonate shunt pathway. (**A**) Intracellular levels of selected non-oxygenated sterols in CD4+ T cells (n = 5). Metabolites include: 7-dehydrocholesterol, 7-dehydrodesmosterol, 8,9-dehydrocholesterol, 14-desmethylanosterol, 24,25-dihydrolanosterol, campesterol, cholestanol, cholestanone, desmosterol, follicular fluid meiosis-activating sterol (FFMAS), lanosterol, sitosterol, zymosterol. Heatmap represented as Log_2_(fold-change) versus vehicle control. (**B**) Oxysterol / sterol ratio in CD4+ T cells (n = 5). (**C**) Changes in enzyme transcript levels within the 24,25-epoxycholesterol shunt pathway, as measured by RNA-seq, in CD4+ T cells (n = 4). (**D**) Overlap between liver X receptor-associated genes and genes differentially-regulated by ABHD11 inhibition in CD4+ T cells (n = 4). All experiments were carried out using human samples. CD4+ T cells were activated with α-CD3 and α-CD28 for 24 h, in the presence and absence of ML-226. Data are expressed as mean, with paired dots representing biological replicates.

**Supplementary Figure 6.**
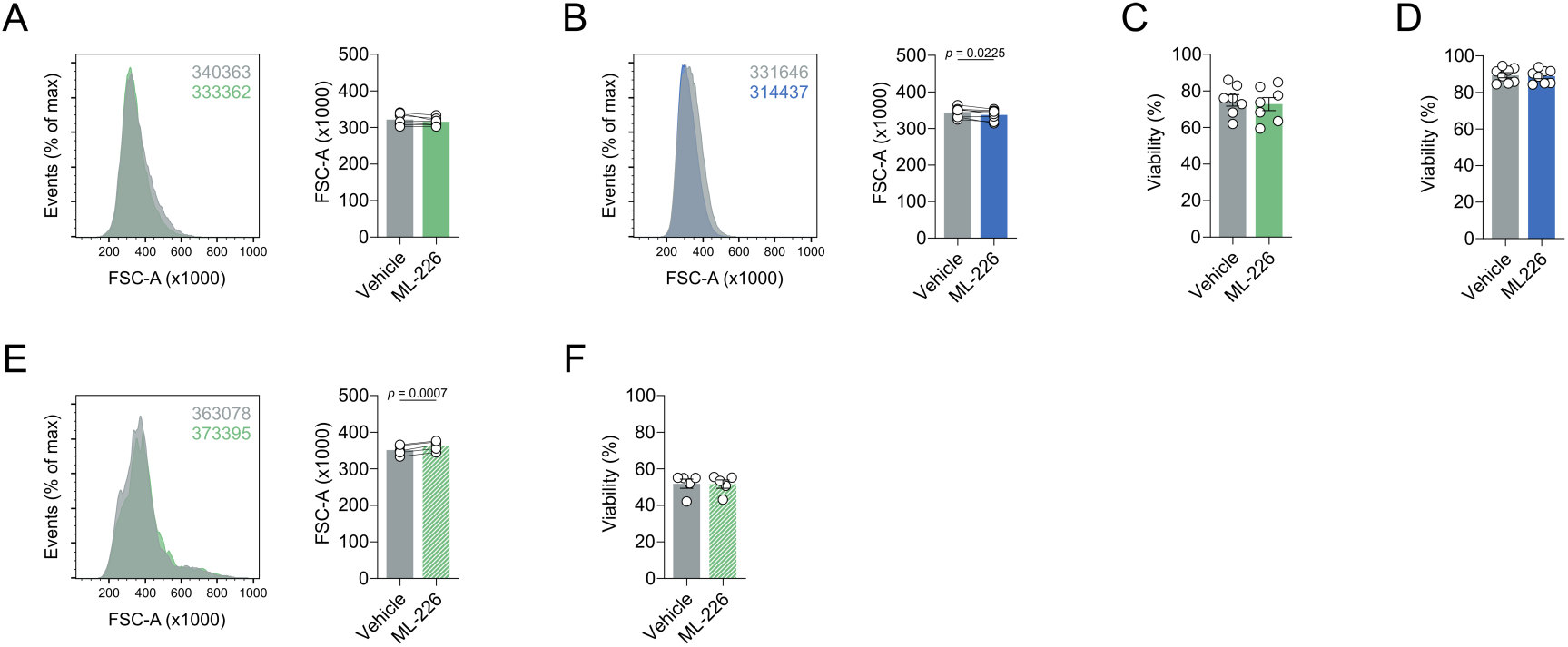
ABHD11 inhibition has no clear effect on T cell size and viability in autoimmune patient cohorts. (**A,B**) Cell size, as determined by forward scatter area, of patient-derived CD4+ T cells in autoimmune cohorts of (**A**) RA (n = 7) and (**B**) T1D (n = 8). (**C,D**) Cell viability, as determined by DRAQ7^®^, in patient-derived CD4+ T cells in autoimmune cohorts of (**C**) RA (n = 7) and (**D**) T1D (n = 8). (**E**) Cell size, as determined by forward scatter area, of patient-derived synovial fluid mononuclear cells (SFMCs; n = 5). (**F**) Cell viability, as determined by DRAQ7^®^, in patient-derived SFMCs (n = 5). All experiments were carried out using human samples. CD4+ T cells were activated with α-CD3 and α-CD28 for 24 h, in the presence and absence of ML-226, unless otherwise stated. Data are expressed as either: mean, with paired dots representing biological replicates; or mean ± SEM.

**Supplementary Figure 7.**
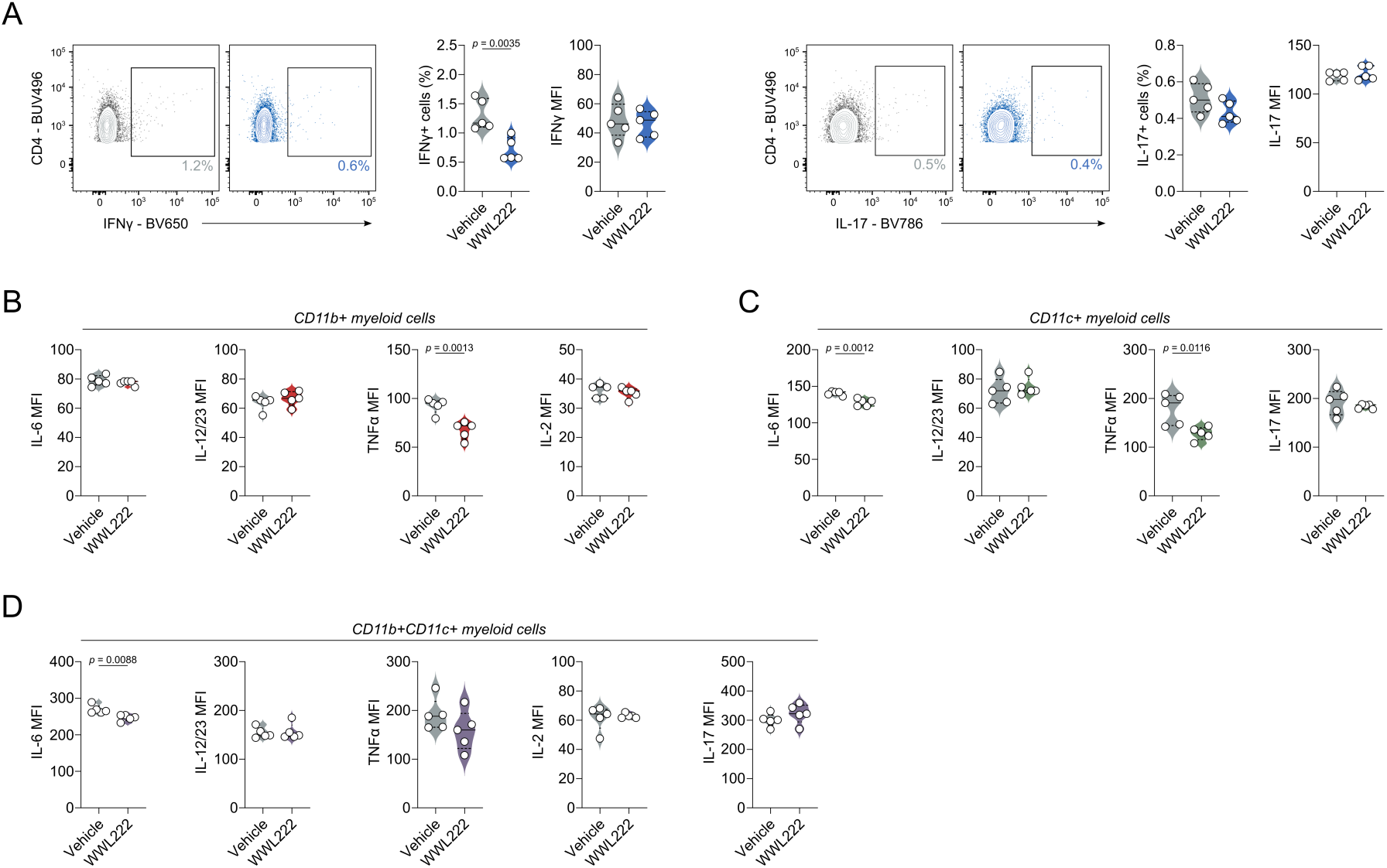
ABHD11 inhibition delays T1D by altering the cytokine profile. (**A**) IFNγ and IL-17 production, as measured by flow cytometry, by CD4+ T cells (n = 5). (**B**) IL-6, IL-12/23, TNFα and IL-2 production, as measured by flow cytometry, by splenic CD11b^+^ myeloid cells (n = 5). (**C**) IL-6, IL-12/23, TNFα and IL-17 production, as measured by flow cytometry, by splenic CD11c^+^ myeloid cells (n = 5). (**D**) IL-6, IL-12/23, TNFα, IL-2 and IL-17 production, as measured by flow cytometry, by splenic CD11b^+^CD11c^+^ myeloid cells (n = 5). All experiments were carried out using murine samples. Mice were injected daily with the indicated dose of WWL222. Data are expressed as median ± interquartile range.

**Supplementary Figure 8.**
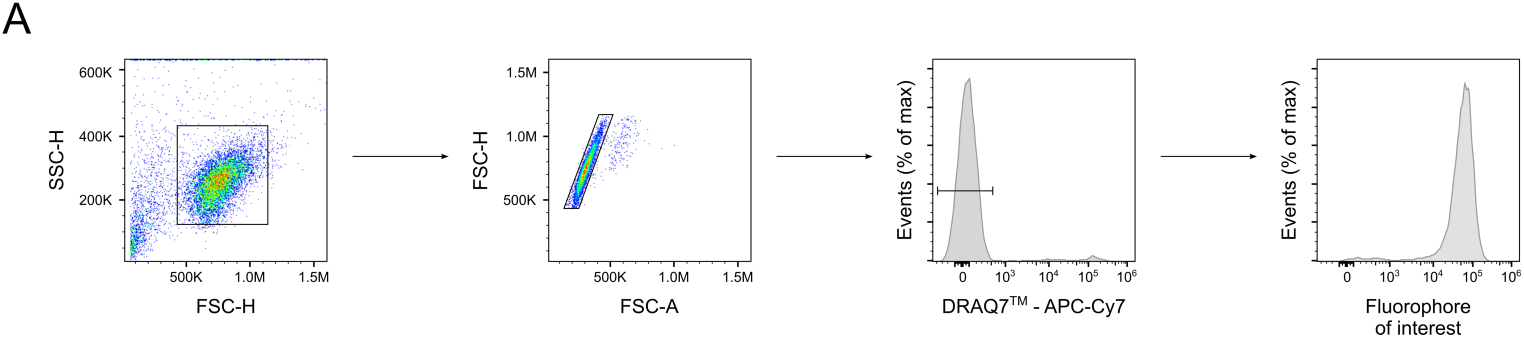
Representative gating strategy. (**A**) Representative gating strategy employed for flow cytometry analysis. Cell doublets were excluded from analysis based on forward scatter-height versus forward scatter-area. Cell death was monitored using DRAQ7™ (1 μM; Biostatus, UK) and dead cells were excluded from analysis.

**Supplementary Table 1.**
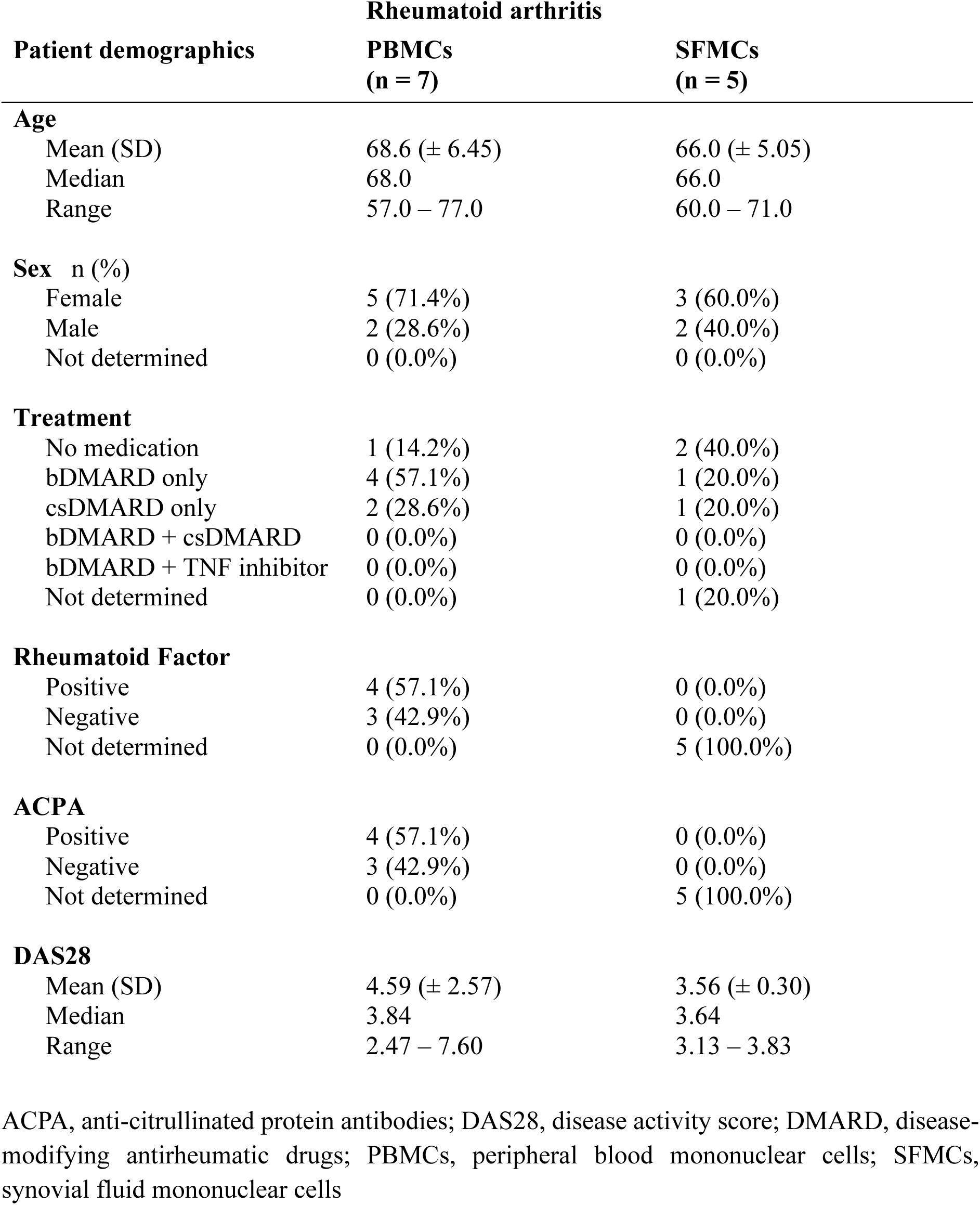
Rheumatoid arthritis patient demographics.

**Supplementary Table 2.** Type 1 diabetes patient demographics.

| Patient demographics | Type 1 diabetes<br>(n = 8) |
| --- | --- |
| **Age** |  |
| Mean (SD) | 36.4 (± 14.3) |
| Median | 32.4 |
| Range | 19.0 – 62.0 |
| **Sex n (%)** |  |
| Female | 7 (87.5%) |
| Male | 1 (12.5%) |
| **Diabetes duration** |  |
| Mean (SD) | 14.2 (± 14.8) |
| Median | 11.06 |
| Range | 1.8 – 47.0 |
| **Treatment** |  |
| Insulin | 8 (100.0%) |

**Supplementary Table 3.**
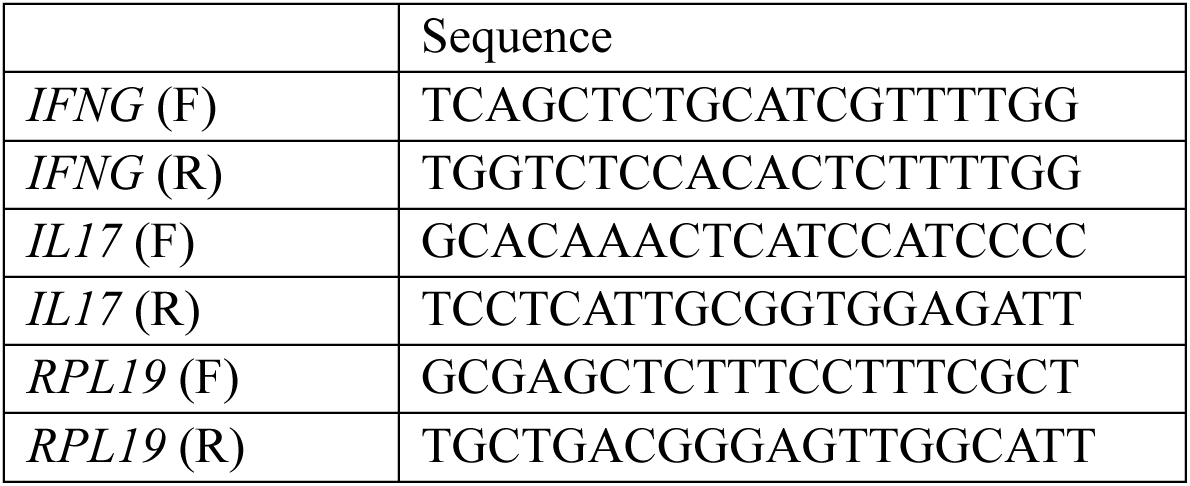
Primer sequences.

